# Evaluating the Nanopore Inosine Model for ADAR-Dependent A-to-I RNA Editing Detection

**DOI:** 10.64898/2026.11.23.768176

**Authors:** Salvatore Di Giorgio, Valerie Griesche

## Abstract

Oxford Nanopore direct RNA sequencing (dRNA-seq) reads native RNA molecules, and its basecaller now includes models that detect RNA modifications directly, among them inosine, the product of adenosine-to-inosine (A-to-I) editing by ADAR enzymes. These models are increasingly used, but how reliably they detect modifications in biological samples has rarely been tested. We evaluated the Dorado inosine model in a mouse macrophage cell line, comparing wild-type cells with ADAR1-knockout cells and in vitro transcribed RNA, before and after an inflammatory stimulus that increases editing. As a reference, we used the editing sites identified by Illumina sequencing of the same cells. We assessed whether the model distinguishes edited from unedited RNA, which negative control is needed to identify editing sites, how many reference sites are recovered, and why sites are missed. The confidence of the Nanopore inosine model alone was not sufficient to identify editing, and a matched ADAR-deficient control was essential. Even with this control, detection was limited by sequencing depth, by low editing levels and by basecalling errors at edited positions. Nanopore sequencing can detect A-to-I editing, but its inosine calls require an ADAR-deficient control and sufficient depth, and should be confirmed by an orthogonal method.

## 1 Introduction

More than one hundred chemical modifications of RNA have been described^1^. Most of them are chemical additions to the base or the ribose, such as methylations, that do not change the identity of the nucleotide. They mainly decorate abundant non-coding RNAs, but a subset occurs on messenger RNA, where they may act as dynamic regulators of gene expression ^2^. The most abundant internal mRNA modification is N6-methyladenosine (m6A), which has been mapped in the transcripts of more than 7,000 human genes^3,4^, followed by marks such as pseudouridine (*ψ*), N1-methyladenosine (m1A), 5-methylcytosine (m5C) and 2*^′^*-O-methylation (Nm) ^5^.

In contrast, RNA editing changes the base itself. In mammals, the predominant form is the deamination of adenosine to inosine (A-to-I) by the ADAR enzymes ^6^; cytidine-to-uridine editing by APOBEC enzymes is far less frequent^7,8^. ADARs act on double-stranded RNA, and most A-to-I editing therefore occurs in non-coding regions, within dsRNA structures formed by inverted repetitive elements, such as Alu elements in primates, B1 and B2 SINEs in rodents, and LINE sequences, in introns and UTRs^9^. Besides recoding a small number of codons, editing modulates RNA structure, splicing and microRNA targeting, and, in innate immunity, marks endogenous dsRNA as “self”, preventing the activation of the antiviral sensors MDA5 and PKR^6,10,11^.

In Illumina-based transcriptomics, A-to-I editing sites are identified as A-to-G mismatches, because reverse transcriptase reads inosine as guanosine ^12^. This approach is the current standard for quantification, but it is indirect: it infers editing from the base pairing of a cDNA copy rather than detecting inosine on the native RNA.

Direct RNA sequencing (dRNA-seq) by Oxford Nanopore Technology (ONT) analyses native RNA molecules: as a strand translocates through a nanopore, its nucleotides, including modified ones, alter the ionic current, so that modifications can in principle be detected from their signal rather than inferred from mismatches ^13,14^. Since the first dRNA-seq chemistry, many computational approaches have been developed to map RNA modifications^15,16^. They fall into three groups: (1) comparative error profiling, which infers modifications from systematic base-calling errors^17–20^; (2) raw-signal analysis, which uses statistical or machine-learning classifiers on current amplitude and dwell time ^21–25^; and (3) modification-aware base-calling, which assigns a modification probability to each base during base-calling, so that modifications can be called at the level of single reads ^16^. These strategies have been benchmarked for specific modifications ^26–28^, but the technology evolves rapidly.

With the SQKRNA004 chemistry, ONT integrated modification-aware models directly into its production base-caller (Dorado), including a model for inosine. Although these models are increasingly used ^15^, independent evaluations remain limited and have focused on m6A and PsU ^29,30^ or on tRNA ^31^; the performance of the inosine model for low-level A-to-I editing in native transcripts has not been assessed. A systematic evaluation is therefore needed to determine whether their convenience comes at the cost of accuracy.

Inosine is a chemically subtle modification that is structurally similar to guanosine ^32^. Deep-learning approaches have improved the discrimination of inosine from guanosine in synthetic or highly edited RNA ^18,33,34^. In cells, however, A-to-I editing is substoichiometric: editing levels rarely exceed 40–50% and average approximately 10% ^35,36^, and many sites lie in AU-rich, repetitive sequences. The nanopore signal at a site is therefore a mixture of few edited and many unedited molecules. Whether a pretrained model can reliably detect such low-level editing in native transcripts, rather than in high-abundance controls, remains an open question.

Here, we evaluated the Dorado inosine model (SQK-RNA004) in a mouse macrophage cell line (RAW 264.7). We compared WT cells with ADAR1-KO cells, which lack ADAR1-dependent editing, and with *in vitro* transcribed (IVT) RNA, which contains no modifications. As a reference, we defined ADAR-dependent editing sites by Illumina sequencing of the same cells. With this design we asked: (i) whether the confidence of the model distinguishes edited from unedited RNA; (ii) which negative control, ADAR1-KO cells or IVT RNA, is needed to identify editing sites; (iii) how many of the reference sites Nanopore recovers and how many of its calls the reference supports; (iv) why reference sites are missed, considering sequencing depth, editing level and the inosine signal in individual reads; and (v) how the basecaller handles edited reads, including possible basecalling errors at the edited position.

## 2 Results

### 2.1 ADAR-dependent A-to-I RNA editing in Illumina sequencing data

To obtain a reference set of editing sites for benchmarking, we characterised A-to-I editing by Illumina RNA sequencing in RAW 264.7 macrophages (immortalised mouse macrophages), comparing wild-type (WT) cells with ADAR1 knockout (ADAR1-KO) cells, unstimulated and after 24 h of IFN-*γ*/LPS stimulation, which modulates editing levels. To restrict the analysis to ADAR-dependent events, candidate sites were selected among positions that were covered by at least 10 reads but showed no A-to-G mismatch in ADAR1-KO cells at either time point. This excludes A-to-G mismatches that arise independently of ADAR, such as genomic variants of the cell line, spontaneous adenosine deamination or sequencing and alignment errors. Positions with A-to-G mismatches in at least two of three wild-type replicates at both time points and with at least five edited reads at a median coverage *≥* 10 were retained^37,38^, yielding 2,971 A-to-I editing sites (**Supplementary Data 1**; Methods, *Illumina RNA-seq and A-to-I editing detection*). Because they were selected against the ADAR1-KO background, these sites are unedited in ADAR1-KO cells by construction (**Fig. S1B–C**).

In wild-type cells, editing levels were low in the absence of stimulation: the median editing frequency across the 2,971 sites was 4%, and only 1,371 sites (46%) were edited in at least 5% of the reads. After 24 h of IFN-*γ*/LPS stimulation, the median frequency rose to 22%, and 2,640 sites (89%) were edited in at least 5% of the reads (**Fig. 1A**), consistent with the induction of the interferon-inducible ADAR1 isoform p150 in activated macrophages ^39–41^. The sites therefore span a broad range of editing levels, which is required to assess the sensitivity of Nanopore detection.

**Figure 1:**
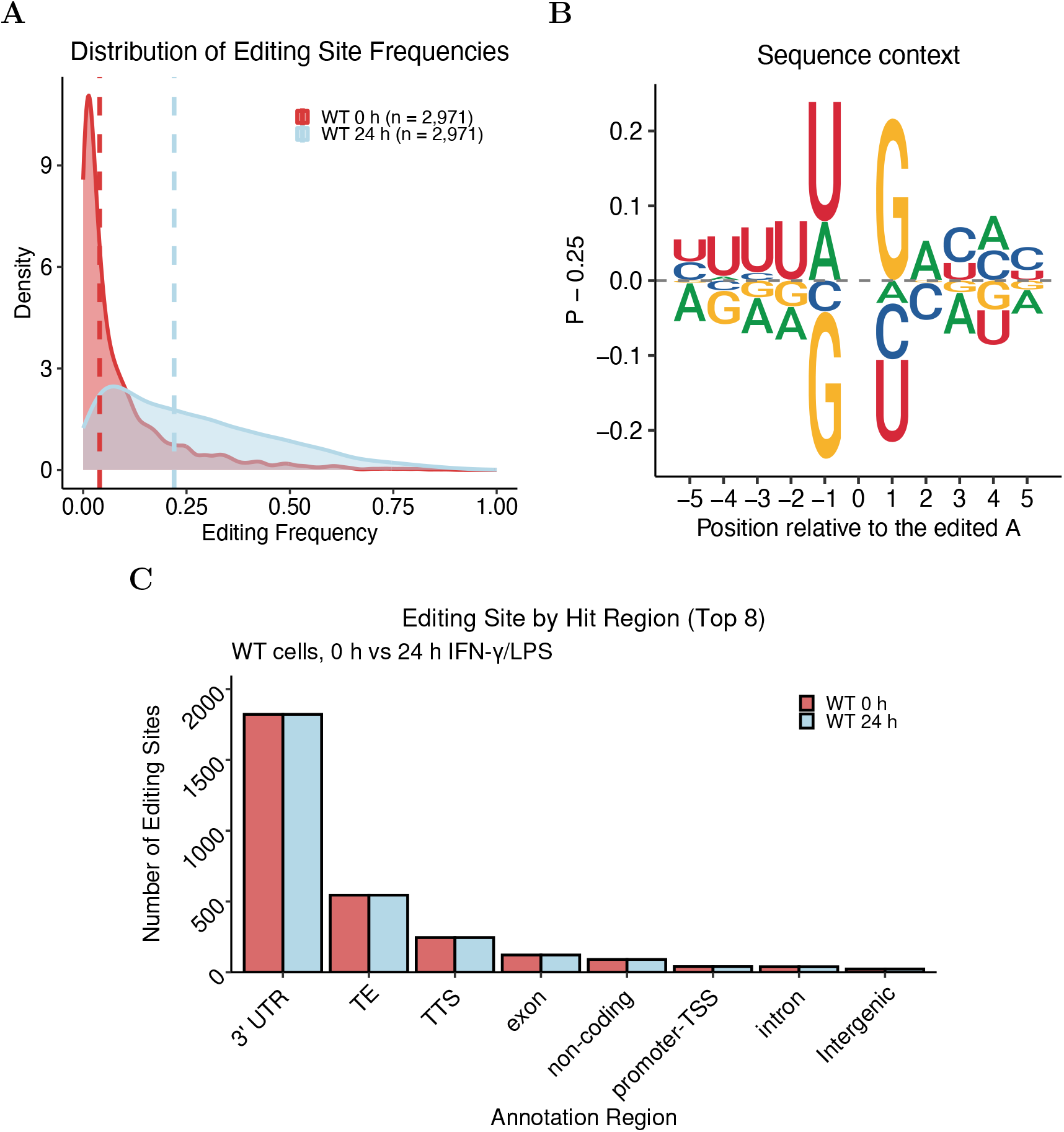
ADAR-dependent A-to-I editing sites identified by Illumina RNA sequencing. Editing sites were defined in wild-type (WT) RAW 264.7 macrophages as A-to-G mismatches absent in ADAR1-KO cells (*n* = 2,971; selection criteria in Methods). Editing frequencies are medians over three replicates. **(A)** Distribution of the editing frequency of the 2,971 sites in WT cells in unstimulated cells (0 h; red) and after 24 h of IFN-*γ*/LPS stimulation (blue); dashed lines, medians (4% and 22%). **(B)** Sequence context of the edited adenosine (*±*5 nt), shown as the frequency of each base minus 0.25; bases above 0 are enriched and bases below 0 are depleted. **(C)** Number of editing sites per genomic region (HOMER annotation; TE, transposable elements: SINE, LINE and LTR; TTS, transcription termination site): 3*^′^* UTR, 1,822 sites (61.3%); TE, 545 (18.3%; SINE, 460). Editing sites and annotation (C) are given in Supplementary Data 1, editing frequencies (A) in Supplementary Data 2.

Several independent features support that the sites reflect genuine ADAR activity. Their sequence context matched the known ADAR preference, with a depletion of guanosine (G) immediately upstream and an enrichment of G downstream of the edited adenosine ^42,43^ (**Fig. 1B**). Most sites were located in 3*^′^* UTRs (61.3%) and a substantial fraction in transposable elements (18.3%), mostly SINEs (15.5%), the preferred substrates of ADAR in double-stranded regions formed by inverted repeats (**Fig. 1C** and **Supplementary Data 1**). None of the sites coincided with single nucleotide polymorphisms (SNPs) of the BALB/c background of RAW 264.7, and only 3.2% (96 sites) coincided with positions that are polymorphic in other mouse strains (Mouse Genomes Project v5, dbSNP build 142; **Fig. S1A**, **Table S2** and **Supplementary Data 2**; Methods, *Annotation and characterisation of editing sites*). Moreover, 46.7% of the sites are listed in the REDIportal catalogue of known editing sites^44,45^ (**Fig. 4B**); the remaining sites are not unexpected, as the mouse REDIportal catalogue is derived from brain and bone marrow and does not include stimulated macrophages. Because sites had to be detected at both time points, editing sites specific to stimulated cells are under-represented in this reference set (Methods, *Stimulation-specific editing sites*). Together, these data define a set of ADAR-dependent A-to-I editing sites spanning diverse genomic contexts and editing levels, which we used as the reference for evaluating Nanopore direct RNA sequencing.

### 2.2 Evaluation of ONT modification-aware base-calling specificity

To assess whether the modification-aware base-calling models can distinguish genuine modifications from background, we analysed the per-read modification probabilities reported by modkit (sample-probs) for m6A, m5C and inosine. WT samples were compared with two types of negative controls: writer knockouts, METTL3-KO for m6A and ADAR1-KO for inosine, in which the respective modification is biologically depleted, and in vitro transcribed (IVT) RNA, which is transcribed from cDNA and therefore contains no modified nucleotides. RNA editing, however, is only partly erased: during reverse transcription, inosine pairs with cytidine and is copied as guanosine, so that IVT RNA carries an unmodified G at A-to-I sites. The modification is lost, but the A-to-G change in sequence is retained. Instead of comparing the shape of the probability distributions, which are normalised within each sample and therefore hide differences in the number of calls, we counted for every probability cutoff the calls above the cutoff per million calls of the respective base (**Fig. 2A-C-E**), and expressed the specificity of each model as the enrichment of these calls in WT relative to the negative control (**Fig. 2B-D-F**; Methods, *Modification probability distributions*). A specific model is expected to yield an enrichment above 1 that increases with the cutoff, whereas an enrichment of 1 indicates that the negative control carries as many confident calls as WT.

**Figure 2:**
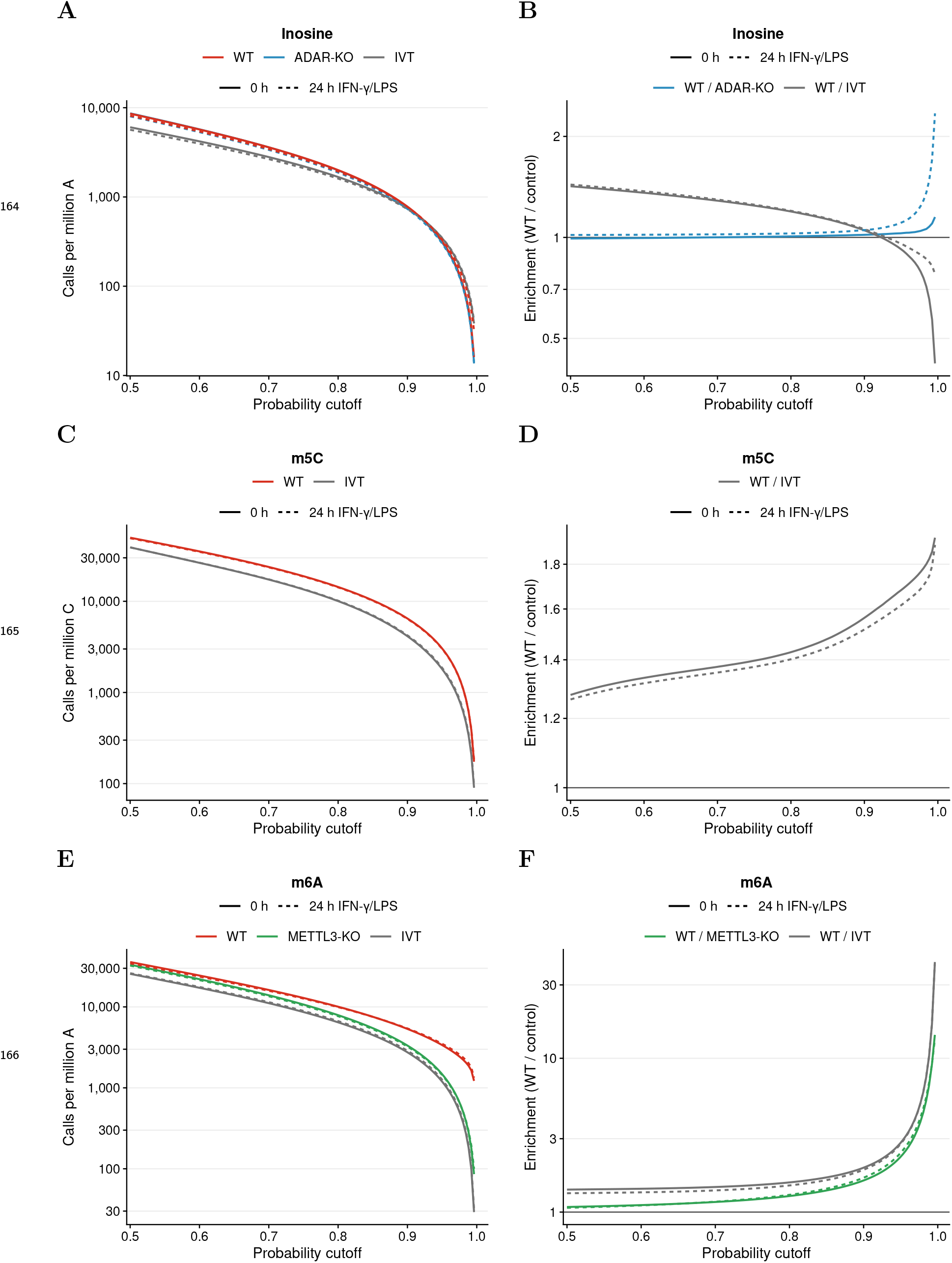
Specificity of modification-aware basecalling for m6A, m5C and inosine. **(A–C-E)** Number of modification calls with probability *≥* cutoff per million calls of the respective base (modkit sample-probs), for wild-type (WT) cells, writer knockouts (METTL3-KO for m6A, ADAR-KO for inosine) and in vitro transcribed RNA (IVT). Lines show the mean over replicates; solid: unstimulated (0 h), dashed: 24 h IFN-*γ*/LPS. **(B–D-F)** Enrichment of calls in WT relative to each negative control. A value of 1 (horizontal line) means the negative control contains as many confident calls as WT. m6A and m5C calls are increasingly enriched in WT at higher cutoffs, whereas inosine calls are equally frequent in WT and ADAR-KO at all cutoffs and less frequent in WT than in IVT at cutoffs > 0.9. Values are given in Supplementary Data 1.

The m6A model behaved as expected for a specific detector. Already without a cutoff, a lower fraction of adenosine calls was classified as m6A in METTL3-KO (3.9%) and IVT (3.0%) than in WT (4.2%) (**Supplementary Data 1**). This difference grew with the confidence of the calls (**Fig. 2E**): at a probability *≥* 0.9, WT carried 5,248 m6A calls per million adenosines compared with 3,236 in METTL3-KO and 2,672 in IVT, and at *≥* 0.99 the difference reached 1,639 versus 179 and 99 calls per million, respectively (**Supplementary Data 1**). Accordingly, the enrichment of m6A calls in WT increased from 1.1-fold (METTL3-KO) and 1.4-fold (IVT) at a cutoff of 0.5 to 9.1-fold and 16.6-fold at 0.99 (**Fig. 2F**). The m5C model also separated WT from the unmodified IVT control, although less strongly, with an enrichment increasing from 1.3-fold at 0.5 to 1.9-fold at 0.99 (**Fig. 2C-D**). For both models the profiles were virtually identical in unstimulated cells and after 24 h of IFN-*γ*/LPS stimulation.

In stark contrast, the inosine model did not distinguish WT from ADAR1-KO cells, in which A-to-I editing is abolished (**Fig. 2A-B**). The fraction of adenosine calls classified as inosine was the same in both genotypes (1.1% in WT and ADAR1-KO), and the number of confident inosine calls was indistinguishable at every cutoff: at a probability *≥* 0.9, WT carried 755 and ADAR1-KO 742 inosine calls per million adenosines (750 versus 712 at 24 h), corresponding to an enrichment of 1.02 (1.05 at 24 h; **Fig. 2B**). Only in the extreme tail (*≥* 0.99), which contained fewer than 60 calls per million adenosines (**Supplementary Data 1**), did the enrichment rise slightly after stimulation (1.10 at 0 h, 1.71 at 24 h). Comparison with IVT RNA revealed an even more striking pattern: although IVT RNA carried fewer inosine calls overall (0.75% of adenosine calls) and at low cutoffs (WT/IVT enrichment of 1.4 at a cutoff of 0.5), the enrichment declined with increasing confidence and dropped below 1 above a probability of about 0.9, reaching 0.57 at 0.99 (0.82 at 24 h). Thus, unmodified IVT RNA contained up to 1.8-fold more high-confidence inosine calls than WT RNA.

Together, these results show that, unlike the m6A and m5C models, the inosine model does not distinguish RNA that contains inosine from RNA that does not when all adenosines are considered: genome-wide, confident inosine calls are as frequent in ADAR1-KO as in WT RNA, increasing the probability cutoff does not enrich them in WT, and IVT RNA, which contains no inosine because edited sites are copied as guanosine, carries even more confident calls than WT RNA. These IVT calls are therefore false positives by definition. Because genuine editing affects only a small fraction of all adenosines, this does not exclude a signal at the edited sites themselves, which we examine below (**Fig. 6**). Consequently, the reported probability is not informative on its own: without a negative control, it cannot be known what fraction of the calls above any cutoff reflects inosine, and a matched ADAR-deficient control is required to identify editing sites from Nanopore data.

### 2.3 ADAR-dependent A-to-I editing in Nanopore direct RNA sequencing data

Since the modification probability alone did not separate edited from unedited RNA (**Fig. 2**) efficiently, we identified candidate inosine sites by differential analysis against a negative control. Because no probability threshold could be derived from the score distributions, the whole analysis was repeated for eight modkit pass thresholds for inosine calls (0.3, 0.5, 0.6, 0.7, 0.8, 0.9, 0.95 and 0.99; canonical adenosine calls at 0.7). For each threshold, per-site inosine levels were compared position by position between WT samples and either ADAR1-KO samples or IVT RNA, separately for unstimulated samples and after 24 h of IFN-*γ*/LPS stimulation (Methods, *Differential inosine analysis*).

In the following, a read was counted as carrying inosine only if the model was at least 80% confident (threshold 0.8). This cut-off recovered the largest share of the Illumina reference sites while keeping the number of unsupported calls low, in particular after stimulation (**Fig. 4C**). With ADAR1-KO cells as negative control, 903,551 positions in unstimulated cells and 1,337,727 positions after 24 h of IFN-*γ*/LPS stimulation were covered well enough to be tested (**Fig. 3A–B**; **Supplementary Data 2**). Only 102 positions differed significantly between unstimulated WT and ADAR1-KO cells (FDR *<* 0.05, *|* log_2_FC*| ≥* 0.5), whereas 2,205 positions did so after stimulation, in line with the stimulation-induced increase in editing observed by Illumina sequencing (**Fig. 1A**). The direction of these changes was consistent with ADAR-dependent editing: all 102 sites at 0 h and 2,174 of 2,205 sites (98.6%) at 24 h showed higher inosine levels in wild-type than in ADAR1-KO cells (**Fig. 3A-B and Supplementary Data 2**). The comparison with IVT RNA yielded more significant positions (1,278 at 0 h and 3,248 at 24 h), but a substantial fraction of them, 287 (22.5%) and 346 (10.7%), showed higher inosine levels in the unmodified IVT RNA than in WT RNA (**Fig. 3C-D and Supplementary Data 2**), which cannot reflect genuine editing. The two negative controls also differed in their dependence on the pileup threshold (**Fig. S2**; **Supplementary Data 2**), although the number of tested positions decreased similarly with increasing threshold for both controls (**Fig. S2A**): with ADAR1-KO, the number of positions with higher inosine levels in wild type was nearly constant across thresholds (96–107 at 0 h and 2,063–2,577 at 24 h, thresholds 0.3–0.99; **Fig. S2B**), while the number of positions in the opposite direction decreased from 90 to 3 at 24 h (**Fig. S2C**). With IVT, in contrast, the number of positions with higher inosine levels in wild type changed by three orders of magnitude, from 22,778 at a threshold of 0.3 to 991 at 0.8 and 27 at 0.99 (0 h; **Fig. S2B**), indicating that most of these differences are driven by low-confidence calls rather than by editing.

**Figure 3:**
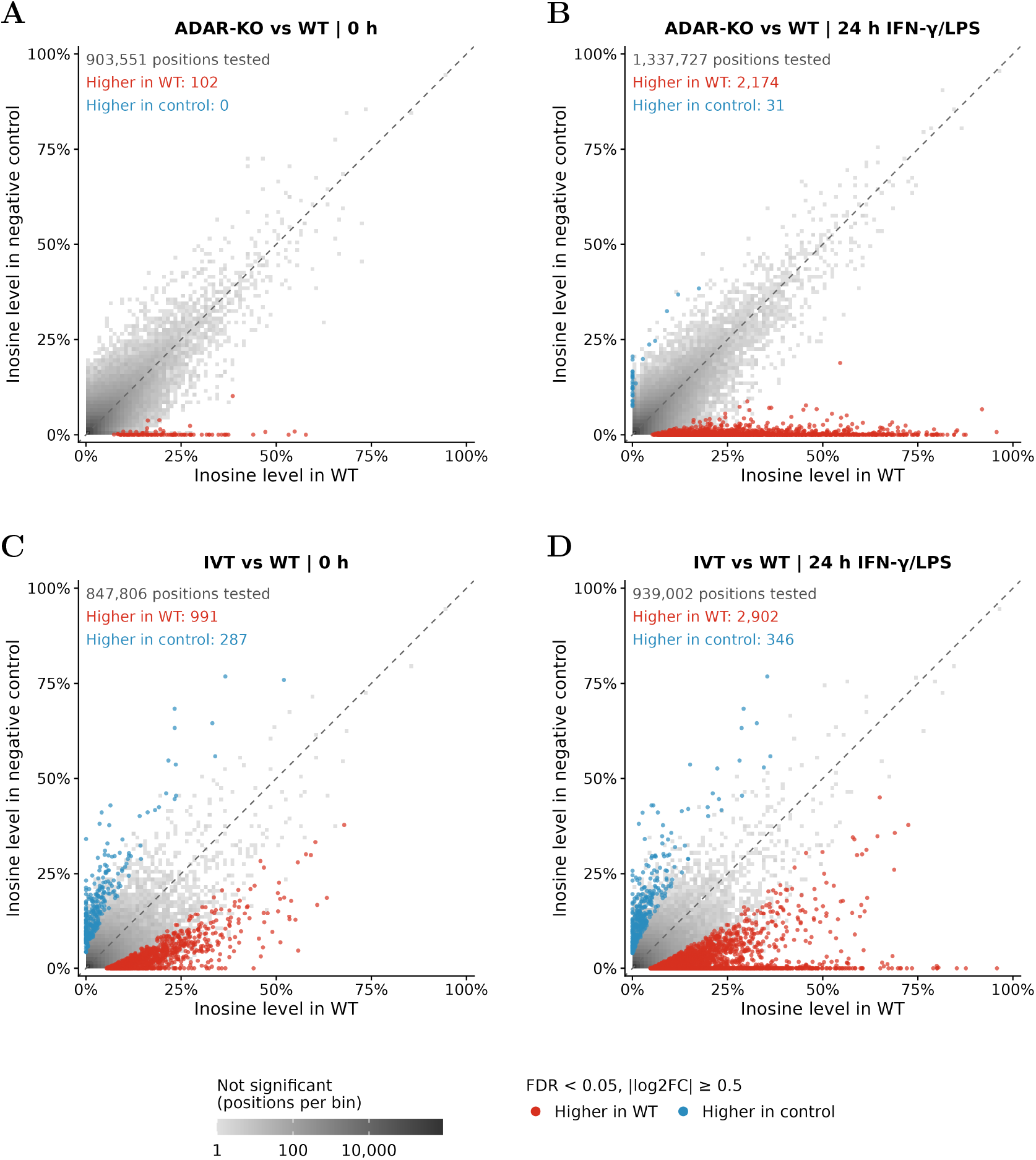
Differential inosine analysis between wild-type cells and negative controls. Mean inosine level per position in WT samples (x axis) and in the negative control (y axis): **(A, B)** ADAR1-KO cells and (C, D) *in vitro* transcribed (IVT) RNA, in unstimulated cells **(A, C)** and after 24 h of IFN-*γ*/LPS stimulation **(B, D)**. Modification calls were obtained with a modkit pass threshold of 0.8 for inosine. Differential modification was tested with the DSS R package; positions with FDR *<* 0.05 and *|* log_2_FC*| ≥* 0.5 are shown as points, red when the inosine level is higher in WT and blue when it is higher in the negative control. Non-significant positions are shown as a grey density (number of positions per bin). The dashed line indicates equal inosine levels. Numbers give the tested positions and the significant positions in each direction. Values are given in Supplementary Data 2.

The positions with higher inosine levels in wild type at either time point were pooled, giving 2,184 positions for ADAR1-KO (102 at 0 h and 2,174 at 24 h; 92 at both) and 3,169 for IVT (991 at 0 h and 2,902 at 24 h; 724 at both). They were retained as candidate sites if they were covered by at least five reads in at least two replicates of at least two sample groups (genotype and time point, i.e. WT and the negative control at 0 h and 24 h), were not modified in the negative control (median frequency *<* 5% at a median coverage *≥* 10) and carried at least five modified reads at a median coverage *≥* 10 in WT (medians over replicates; Methods, *Inosine site calling*). The replicate requirement removed no position; 29 positions were removed because they were modified in ADAR1-KO and 324 because they carried fewer than five modified reads in WT. This yielded 1,831 candidate editing sites with ADAR1-KO (83.8% of the pooled positions) and 2,251 with IVT (71.0%; 538 positions modified in IVT and 380 with fewer than five modified reads) as negative control (**Supplementary Data 2**). Across thresholds, the number of candidate sites changed little with ADAR1-KO (2,195 at 0.3 to 1,090 at 0.99) but about 30-fold with IVT (20,826 to 664; **Fig. S2D**). Consistent with the differential analysis, the ADAR1-KO-derived sites were largely induced by stimulation: their median editing frequency in wild-type cells rose from 0% at 0 h, when only 309 of them were edited in at least 5% of the reads (the minimum frequency used for the comparison with Illumina), to 21% at 24 h, when all but one were (**Fig. S3A**; compare **Fig. 1A**). At the sites detected by both platforms, Nanopore and Illumina editing frequencies were correlated (Spearman *ρ* = 0.63 at 0 h and 0.60 at 24 h), with Nanopore yielding lower frequencies mainly at low editing levels (**Fig. S3B**). As these sites had to pass the Nanopore calling criteria, this correlation is descriptive; the agreement of the editing levels over all covered Illumina sites is examined below (**Fig. 7B**).

We next compared the ONT candidates with the Illumina editing sites (**Fig. 4A**). Of the 1,831 ADAR1-KO-derived sites, 716 (39.1%) coincided with an Illumina editing site at the identical position (Methods, *Overlap between Nanopore and Illumina editing sites*), compared with only 446 of the 2,251 IVT-derived sites (19.8%). The same difference was seen against the REDIportal catalogue of known editing sites ^44,45^, which contained 42.4% of the ADAR1-KO-derived sites, a proportion close to that of the Illumina sites (46.7%), but only 20.2% of the IVT-derived sites (**Fig. 4B**). To quantify detection performance while accounting for the lower Nanopore sequencing depth (see Section 2.4), we restricted the comparison to positions covered by at least 10 reads in both platforms and in the respective Nanopore negative control (Methods, *Benchmarking of Nanopore inosine calls against Illumina*). We report precision (the fraction of Nanopore sites that are also Illumina sites), sensitivity (the fraction of Illumina sites that are also detected by Nanopore) and the *F*_1_ score (their harmonic mean). Under these conditions, the ADAR1-KO-based approach reached a precision of 60.3% and a sensitivity of 21.4% at 0 h (143 of 667 Illumina sites recovered; *F*_1_ = 31.6%), and of 43.4% and 37.2% at 24 h (705 of 1,896; *F*_1_ = 40.1%; **Fig. 4C**; **Table S4**; **Supplementary Data 2**). Expressed symmetrically, as the Jaccard index, 18.8% (0 h) and 25.1% (24 h) of all sites detected by either platform were detected by both (**Table S4**). These values were stable when the minimum read depth was raised from 10 to 60 reads (precision 57–62% at 0 h and 43–45% at 24 h; **Fig. S4A**), and against the REDIportal catalogue the precision of the ADAR1-KO-based sites was of the same order as that of the Illumina sites themselves (83.1% versus 75.8% at 0 h and 46.0% versus 51.3% at 24 h; **Fig. S4C**; Methods, *Benchmarking against REDIportal*).

**Figure 4:**
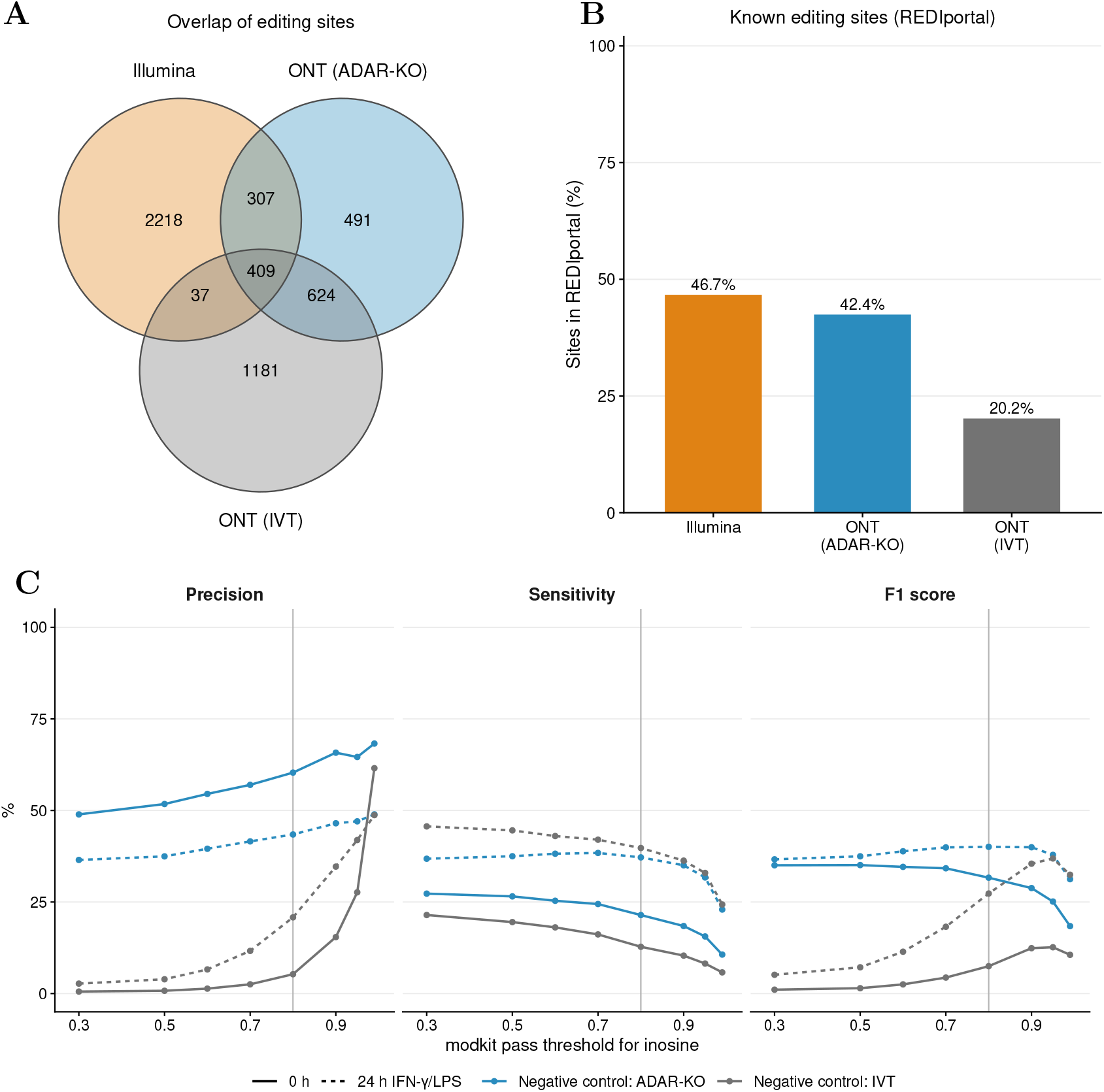
Comparison of Nanopore inosine sites with Illumina editing sites. **(A)** Overlap between the Illumina editing sites and the Nanopore (ONT) candidate editing sites obtained with ADAR1-KO or IVT samples as negative control (modkit pass threshold 0.8). **(B)** Fraction of the Illumina editing sites and of the Nanopore candidate sites contained in the REDIportal catalogue of known A-to-I editing sites. **(C)** Precision, sensitivity and *F*_1_ score of the Nanopore calls relative to the Illumina editing sites across modkit pass thresholds for inosine, using ADAR1-KO (blue) or IVT RNA (grey) as negative control, in unstimulated cells (0 h, solid lines) and after 24 h of IFN-*γ*/LPS stimulation (dashed lines). The vertical line marks the threshold used in A and B. In C, only positions covered by at least 10 reads in both datasets. In C, only positions covered by at least 10 reads in both platforms (and, for Nanopore, in the respective negative control) were evaluated, and calls were matched one-to-one at the identical position. REDIportal (mouse, derived from brain and bone marrow) does not include macrophage- or stimulation-specific sites. Values are given in Supplementary Data 2.

The IVT-based approach performed markedly worse, with a precision of 5.3% at 0 h and 20.8% at 24 h (*F*_1_ = 7.5% and 27.3%), i.e. 80–95% of its calls were not supported by Illumina sequencing, independently of the minimum read depth (**Fig. S4B**) and also when REDIportal was used as reference (precision 7.9% and 21.2%; **Fig. S4C**). Across pileup thresholds, increasing the threshold raised precision at the cost of sensitivity for both approaches. The ADAR1-KO-based approach had a higher precision than the IVT-based approach at every threshold and a higher *F*_1_ score at every threshold except 0.99 after stimulation, where both converged (*F*_1_ 31.2% versus 32.4%, precision 48.9% versus 48.7%). Its *F*_1_ score was highest at 0.8 after stimulation (40.1%; 39.9–40.0% at 0.7 and 0.9) and at 0.3–0.5 in unstimulated cells (35.0–35.1%; **Fig. 4C**). Because the IVT samples were less deeply sequenced, the two controls were evaluated on different numbers of Illumina sites (e.g. 667 versus 415 at 0 h). When both were scored on the same sites, covered by at least 10 reads in the WT, ADAR1-KO and IVT samples, the result was unchanged: at threshold 0.8, the ADAR1-KO-based calls reached a precision of 57.6% versus 5.2% at 0 h and 41.4% versus 21.4% at 24 h, and they had the higher *F*_1_ score at every threshold (**Fig. S5**).

The Illumina sites were used as a reference set, not as a ground truth: these metrics measure how many of the reference sites the Nanopore inosine calls recover and how many of the calls are supported by the reference, not the absolute validity of a site. Sites called only by Nanopore (false positives, FP, in this comparison) or present only in the reference (false negatives, FN) are therefore not necessarily incorrect: they may represent genuine editing sites that are detectable by only one technology, for example in repetitive regions where short reads map ambiguously, or platform-specific artifacts. Consistent with this, 285 of the 1,115 Nanopore-only sites obtained with ADAR1-KO (25.6%) and 895 of the 2,255 Illumina-only sites (39.7%) are known editing sites in REDIportal, compared with 68.7% of the reference sites recovered by Nanopore and only 9.2% of the Nanopore-only sites obtained with IVT (**Supplementary Data 2**). Moreover, because the Illumina reference required editing at both time points, it did not include positions edited only after stimulation; 194 of the 1,115 Nanopore-only sites (17.4%) were such positions, showing A-to-G mismatches in at least two of three WT replicates at 24 h but not at 0 h, and they were more often listed in REDIportal than the remaining Nanopore-only sites (30.9% versus 24.4%; **Supplementary Data 2**; Methods, *Stimulation-specific editing sites*). When these stimulation-specific positions were added to the Illumina reference, the precision of the ADAR1-KO-based approach at 24 h rose from 43.4% to 54.8%, whereas its sensitivity decreased from 37.2% to 22.2%, because Nanopore recovered only 8.8% of the added positions (**Table S3**; **Supplementary Data 2**). These positions rest on two or three replicates at a single time point and are mostly edited at low levels, so that they are a less reliable reference, which is why they were not included in the main benchmark. The Nanopore-only sites also differed in their genomic context (Methods, *Annotation and characterisation of editing sites*): 141 of them (12.6%) were located on unplaced genomic contigs, where no Illumina editing site was found (**Fig. S6B**), and they were enriched in SINE B2 elements (11.0% versus 5.1% of the Illumina-only and 2.5% of the shared sites), whereas the Illumina-only sites were enriched in SINE B1 (Alu-like) elements (12.1%) (**Fig. S6C**). All three classes showed the ADAR sequence preference (depletion of G at position *−*1 and enrichment of G at position +1), although less pronounced for the Nanopore-only sites, which are therefore likely to contain both genuine editing sites in regions poorly accessible to short reads and false-positive calls (**Fig. S6A**). Conversely, absence from REDIportal does not imply that a site is false: the mouse REDIportal catalogue is derived from brain and bone marrow, so editing sites that are specific to macrophages or induced by IFN-*γ*/LPS stimulation are not expected to be represented, which likely also explains why only about half of the Illumina sites (46.7%) are listed. Nanopore calls outside the Illumina reference, and reference sites missed by Nanopore, will therefore require orthogonal validation to distinguish editing events detectable by a single technology from false positives.

Thus, a matched ADAR-deficient control is required to obtain a set of Nanopore inosine calls that is enriched for genuine A-to-I editing. Even with this control, however, Nanopore recovered at most 37% of the Illumina editing sites that were sufficiently covered by both technologies, and 40–57% of its calls were not confirmed by Illumina.

### 2.4 Depth and editing level limit the detection of editing sites by Nanopore

To understand why Nanopore recovered only a minority of the Illumina editing sites, we compared the read depth of both platforms at the benchmarked positions (Methods, *Read depth at the benchmark sites*). At the 2,971 Illumina editing sites, the median wild-type depth was 114 (0 h) and 99 (24 h) reads for Illumina but only 37 and 42 reads for Nanopore, and 99.5% of the sites, compared with 73.8% and 75.5%, were covered by at least 10 reads (median depth per sample 83–122 for Illumina and 22–90 for Nanopore; **Fig. S7**; **Supplementary Data 2**). The IVT samples were the least covered (median 6 and 16 reads; 43.3% and 59.6% of the sites with *≥* 10 reads), which further reduces the number of positions that can be evaluated with this control.

Depth differed markedly between the shared sites and sites present only in the reference or only in the Nanopore calls (ADAR1-KO-based Nanopore sites; classes as in the Venn diagram of **Fig. 4A**) (**Fig. 5A**). Shared sites were well covered by Nanopore (median 72 and 78 reads; *≥* 96% of the sites with *≥* 10 reads), whereas Illumina-only sites had a median Nanopore depth of only 25 reads at both time points, with about one third of them (33.3% and 32.3%) below 10 reads, although their Illumina depth was similar to that of the shared sites (median 113 and 97 reads). Conversely, Nanopore-only sites had a lower Illumina depth (median 26 reads at 0 h and 49 at 24 h; 67.9% and 82.1% with *≥* 10 reads). Thus, part of the sites found in only one dataset reflects insufficient depth in the other dataset rather than a true difference in detection. Nanopore depth at individual sites followed the Illumina depth only in part (Spearman *ρ* = 0.70 at 0 h and 0.57 at 24 h; **Fig. 5B**), so a high coverage in the reference did not guarantee that a site could be evaluated in the Nanopore data. This is consistent with the coverage profile of direct RNA sequencing, which starts at the poly(A) tail and decreases towards the 5*^′^* end of the transcript.

**Figure 5:**
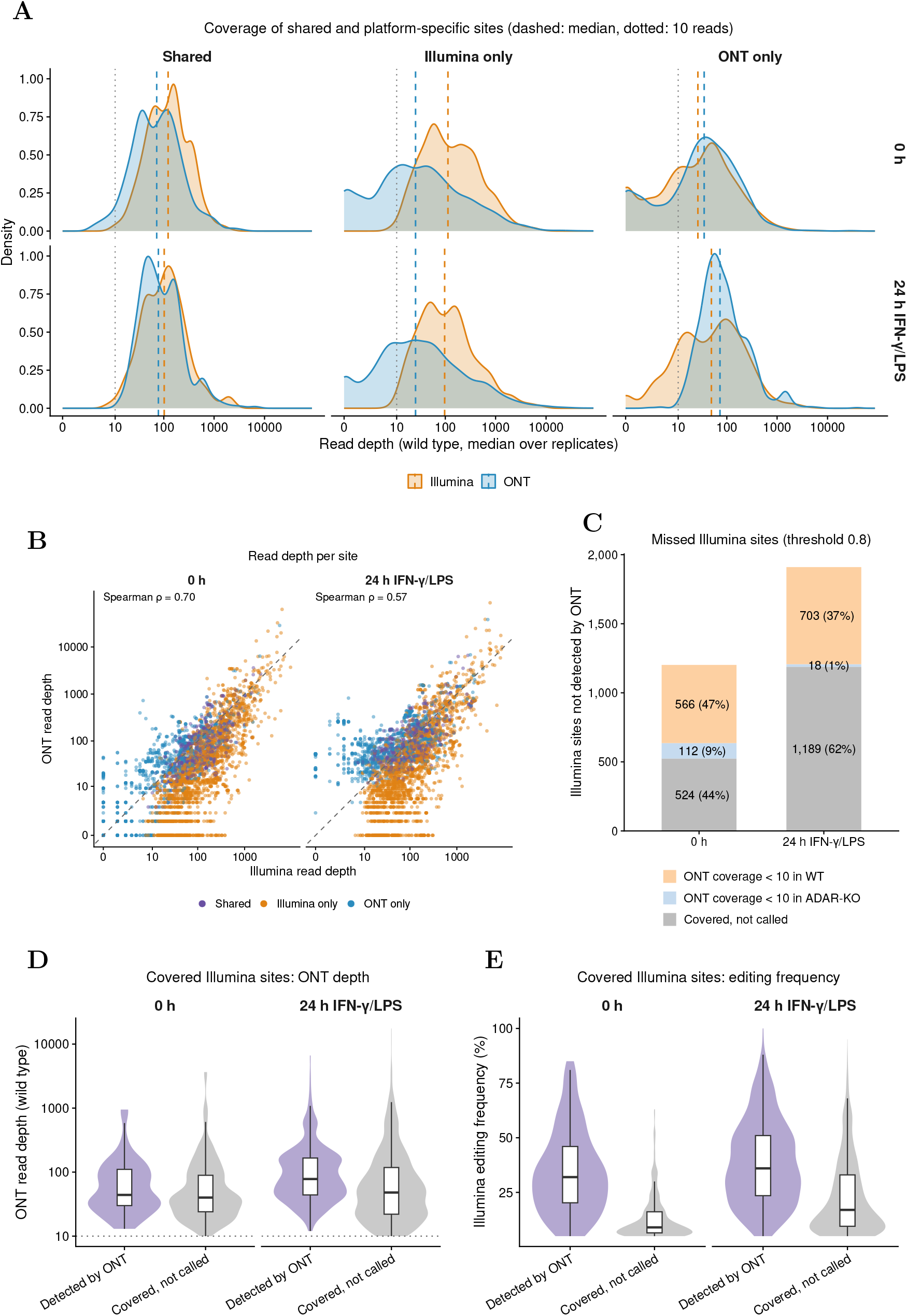
Depth and editing level limit the detection of Illumina editing sites by Nanopore. Editing sites were classified by identical genomic position as detected by both platforms (Shared, *n* = 716), by Illumina only (*n* = 2,255) or by Nanopore only (*n* = 1,115; ADAR1-KO-based candidate sites, pileup threshold 0.8). Read depth was computed with `samtools depth` (MAPQ *≥* 25) and summarised as the median over the wild-type replicates. **(A)** Distribution of Illumina (orange) and Nanopore (blue) read depth for each site class in WT unstimulated cells (0 h) and after 24 h of IFN-*γ*/LPS stimulation. Dashed lines, median depth; dotted line, 10 reads (the minimum coverage used for benchmarking). The x axis is on a log_10_(depth + 1) scale. **(B)** Illumina versus Nanopore read depth per site, coloured by site class. Dashed line, identity; *ρ*, Spearman correlation coefficient. **(C)** Illumina editing sites (detected in at least two replicates with coverage *≥* 10 and editing frequency *≥* 5%; 1,348 at 0 h and 2,617 at 24 h) that were not detected by Nanopore, classified by the first criterion that excludes them: Nanopore coverage *<* 10 reads in wild type, Nanopore coverage *<* 10 reads in the ADAR1-KO control, or covered in both but not called. Numbers give sites and the percentage of missed sites. **(D, E)** Illumina editing sites detected by Nanopore compared with those covered by *≥* 10 reads in both Nanopore samples but not called (grey in C): **(D)** Nanopore wild-type read depth (log scale; dotted line, 10 reads) and **(E)** Illumina editing frequency (median over replicates). Violins show the distribution, boxes the median and interquartile range; whiskers extend to 1.5*×* the interquartile range. Here, an Illumina site counts as detected if a Nanopore candidate site lies at the same position, without the strand, read-depth and one-to-one matching rules of the benchmark; the numbers of detected sites (146 at 0 h and 707 at 24 h) therefore differ slightly from the true positives in **Table S4** (143 and 705). Values are given in Supplementary Data 2.

We then asked how many of the Illumina editing sites missed by Nanopore could be explained by depth (**Fig. 5C**). Of the 1,202 Illumina sites not detected by Nanopore at 0 h, 566 (47.1%) had fewer than 10 Nanopore reads in wild type and 112 (9.3%) in the ADAR1-KO control, so that more than half could not be evaluated; 524 (43.6%) were sufficiently covered but not called. After stimulation, the picture changed: of 1,910 missed sites, 703 (36.8%) and 18 (0.9%) lacked Nanopore depth in WT or ADAR1-KO, respectively, whereas the majority, 1,189 (62.3%), were covered but not called. Insufficient depth therefore accounts for about half of the missed sites in unstimulated cells but only for about a third after stimulation.

A depth of 10 reads, however, does not guarantee that an editing event can be detected, since the calling requires at least five modified reads and a frequency of at least 5%. We therefore compared the covered but missed Illumina sites with those detected by Nanopore (**Fig. 5D-E**). In unstimulated cells, both groups had a similar Nanopore depth (median 40 and 44 reads; difference of medians *−*4 reads), but the missed sites were edited at a much lower frequency according to Illumina (median 9% versus 32%; difference *−*23 percentage points), and half of them (51.3%) were edited in fewer than 10% of the reads. After stimulation, the missed sites had both a lower depth (median 48 versus 78 reads; difference *−*30 reads) and a lower editing frequency (median 17% versus 36%; difference *−*19 percentage points; **Supplementary Data 2**). Combining both quantities, the expected number of edited Nanopore reads (Nanopore depth *×* Illumina editing frequency) was below five for 50.4% of the missed sites at 0 h and 27.3% at 24 h, compared with 15.8% and 2.4% of the detected sites. Thus, most Illumina sites missed by Nanopore are either insufficiently covered or edited at a frequency too low to yield enough modified reads at the available depth, and the stimulation-induced increase in editing frequency (**Fig. 1A**) likely contributes to the higher sensitivity of Nanopore after stimulation.

### 2.5 Nanopore detects an inosine signal at most missed sites, but too rarely to call it

To test whether the Illumina sites that were sufficiently covered but not called by Nanopore were invisible to the inosine model or carried a signal too weak to be called, we extracted the per-read modification probabilities at these sites and at the sites detected by Nanopore (*±*5 nt) in WT, ADAR1-KO and IVT samples, and applied to every read the same inosine threshold as in the pileups used for site calling (*≥* 0.8; **Fig. 6**; Methods, *Per-read inosine probabilities at detected and missed sites*). At the detected sites, a median of 11.2% (0 h) and 25.4% (24 h) of the WT reads carried an inosine call, compared with 0% in ADAR1-KO cells and IVT RNA (**Fig. 6A**; **Supplementary Data 2**). At the missed sites the signal was weaker but present: the median fraction of inosine calls in WT was 1.4% at 0 h and 4.6% at 24 h, again 0% in both controls, and it exceeded the ADAR1-KO level at 69.6% (364 of 523) and 91.6% (1,085 of 1,185) of these sites (median difference between WT and ADAR1-KO at the same site 1.2 and 4.3 percentage points). However, only 10.5% (0 h) and 46.2% (24 h) of them reached the 5% frequency required for calling. Accordingly, reads with an inosine probability *≥* 0.8 were about eight times more frequent in WT than in ADAR1-KO cells at the missed sites (1.3% versus 0.2% at 0 h and 2.8% versus 0.4% at 24 h), but far less frequent than at the detected sites (10.0% and 16.4%; **Fig. 6D**).

**Figure 6:**
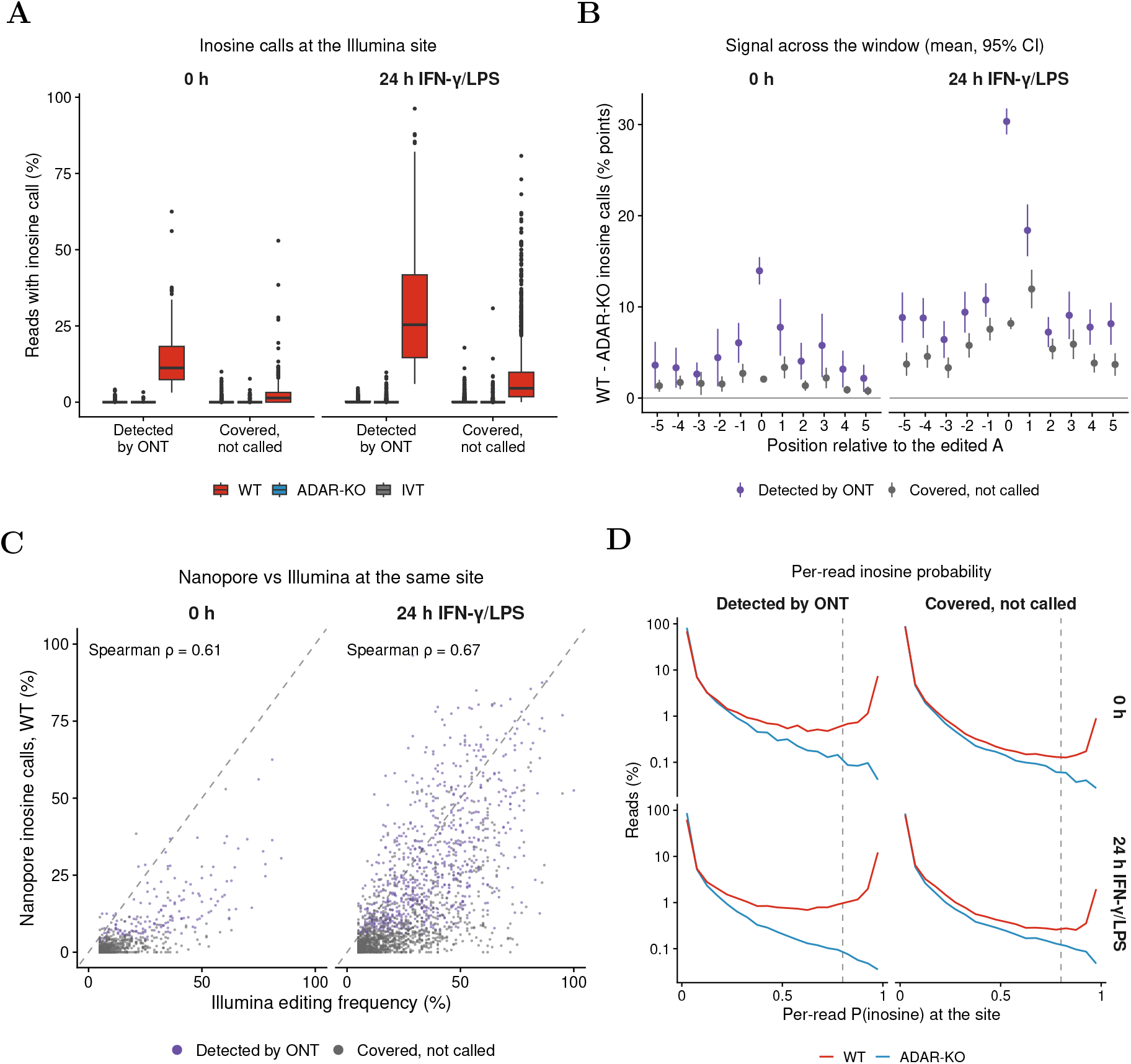
Per-read inosine signal at Illumina editing sites detected or missed by Nanopore. Illumina editing sites detected by Nanopore (purple) or covered by *≥* 10 Nanopore reads in WT and ADAR1-KO samples but not called (grey; ADAR1-KO-based sites, modkit pass threshold 0.8). Per-read probabilities were extracted with `modkit extract full`, calls were made per read with the pileup thresholds (inosine 0.8, other classes 0.7), and reads of the replicates were pooled (sites with *≥* 10 passing reads). **(A)** Percentage of reads with an inosine call at the edited adenosine in WT (red), ADAR1-KO (blue) and IVT (grey) samples. **(B)** Excess of inosine calls in WT over ADAR1-KO along the *±*5 nt window. For each site and each adenosine of the window (x axis; 0, edited adenosine; negative, upstream; positive, downstream), the percentage of reads with an inosine call in ADAR1-KO was subtracted from that in WT; the y axis shows this difference in percentage points (0, no difference), as mean and 95% confidence interval over sites. **(C)** Percentage of inosine calls in WT Nanopore reads versus the Illumina editing frequency of the same site. Dashed line, identity; *ρ*, Spearman correlation. **(D)** Distribution of the per-read inosine probability at the edited adenosine in WT and ADAR1-KO samples (log_10_ scale); dashed line, pass threshold (0.8). In all panels, unstimulated cells (0 h) and cells stimulated for 24 h with IFN-*γ*/LPS are shown separately. Values are given in Supplementary Data 2.

Per site, the fraction of inosine calls in WT correlated with the Illumina editing frequency (Spearman *ρ* = 0.61 at 0 h and 0.67 at 24 h), with the missed sites at the low end of both distributions (**Fig. 6C**). The difference between WT and ADAR1-KO was largest at the edited adenosine but was also elevated at neighbouring adenosines, most strongly at position +1 (**Fig. 6B**), consistent with a partial shift of the inosine signal to the adjacent base. Thus, at most Illumina sites that Nanopore covered but did not call, the inosine model does detect editing; the signal is lost because few edited reads pass the probability threshold at the available depth, so that the site falls below the frequency and modified-read criteria of the calling.

### 2.6 About half of the inosine signal is basecalled as guanosine

The Dorado modification model assigns an inosine probability only to bases that the basecaller has called A. Because inosine is structurally similar to guanosine, edited reads may instead be basecalled G; such reads appear only as an A-to-G mismatch and are counted by the modkit pileup neither as modified nor in the valid coverage. We therefore counted, per position and strand, the reads basecalled G at a reference adenosine in all Nanopore samples (Methods, *Inosine reads basecalled as guanosine*). At 10,000 random adenosines, 0.12–0.22% of the reads were basecalled G in every sample type, the background basecalling error (**Fig. 7A**; **Supplementary Data 2**). At the Illumina editing sites, G calls were twice as frequent in WT as in ADAR1-KO cells at 0 h (0.8% versus 0.4%) and thirteen times as frequent after stimulation (5.1% versus 0.4%), following editing. IVT RNA, which is transcribed from cDNA of the wild-type RNA, carried even more G calls at these sites (1.5% and 8.5%), as reverse transcription copies inosine as guanosine; at edited positions, IVT RNA is therefore not an adenosine-only control. Of the edited reads in WT (excess over ADAR1-KO), 51% (0 h) and 56% (24 h) were basecalled G rather than called as inosine. This fraction depended on the sequence context, from 16% for 5*^′^*-CAC-3*^′^* to 71% for 5*^′^*-UAG-3*^′^*, the preferred ADAR context and the most frequent context among the Illumina sites (**Fig. S8A**).

**Figure 7:**
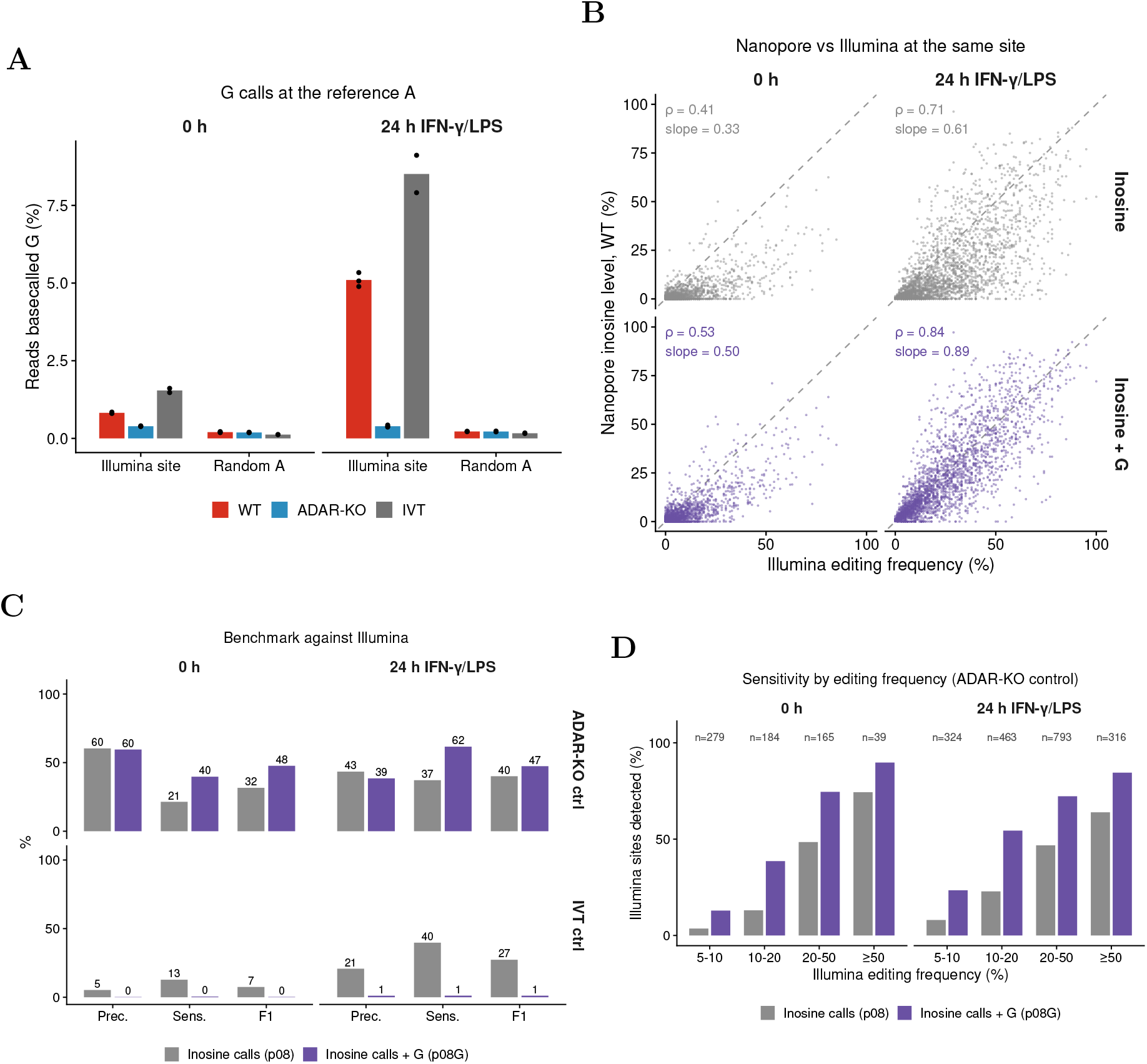
About half of the inosine signal is basecalled as guanosine. Reads basecalled G at a reference adenosine were counted per position and strand with `samtools mpileup` in all Nanopore samples; replicates were pooled per genotype and time point. **(A)** Percentage of reads basecalled G at the Illumina editing sites and at 10,000 random adenosines in WT (red), ADAR1-KO (blue) and IVT (grey) samples (sites with *≥* 10 reads called A or G); bars, mean; points, replicates. **(B)** Nanopore editing level in WT versus the Illumina editing frequency of the same site, computed from inosine calls only (pileup threshold 0.8; top) or from inosine calls plus reads basecalled G (bottom); sites with *≥* 10 Nanopore reads and Illumina sites detected in at least two replicates. Dashed line, identity; *ρ*, Spearman correlation; slope of a linear fit through the origin. **(C)** Precision, sensitivity and *F*_1_ score of the Nanopore candidate sites relative to the Illumina editing sites, with inosine calls only (grey) or with the reads basecalled G counted as modified in all samples, controls included (purple), for the ADAR1-KO (top) and IVT (bottom) control; benchmark as in Fig. 4C (coverage *≥* 10, frequency *≥* 5%, identical position). **(D)** Percentage of the Illumina editing sites detected by Nanopore (ADAR1-KO control) by Illumina editing frequency; *n*, Illumina sites per class. Values are given in Supplementary Data 2. See Methods, *Inosine reads basecalled as guanosine*.

Counting these reads as modified (inosine calls + G; reads basecalled G added to the modified reads and to the coverage) brought the Nanopore editing level close to the Illumina editing frequency after stimulation (Spearman *ρ* from 0.71 to 0.84; slope from 0.61 to 0.89), and improved the agreement also at 0 h (*ρ* from 0.41 to 0.53; slope from 0.33 to 0.50; **Fig. 7B**); because noise in the Illumina frequencies flattens such slopes, agreement was also assessed directly (see below). At 24 h, the Nanopore level then reached 86–93% of the Illumina level in all frequency classes, compared with 48–51% with inosine calls alone; at 0 h it remained lower (28–56% versus 16– 33%; **Fig. S8B**). Measures of agreement, rather than association, gave the same picture: Lin’s concordance correlation coefficient (CCC)^46^, which measures how closely the points lie on the identity line, increased from 0.48 to 0.66 at 0 h and from 0.60 to 0.81 at 24 h, and the mean difference between the Nanopore and the Illumina level (bias) decreased from *−*5.6 to *−*4.0 and from *−*10.4 to *−*2.1 percentage points; the limits of agreement remained wide (about *±*25 percentage points at 24 h), so that the level of an individual site can still deviate considerably (**Fig. S8C**).

We then repeated the complete genome-wide analysis, differential test and site calling, with the G reads counted, at the pass threshold of 0.8. With the ADAR1-KO control, the number of positions significantly more modified in WT increased from 102 to 171 at 0 h and from 2,174 to 3,927 at 24 h, although 71% more positions were tested (2,284,725 versus 1,337,727 at 24 h), and the number of candidate sites increased from 1,831 to 3,408 (**Supplementary Data 2**). Against the Illumina sites, sensitivity increased from 21.4% to 39.7% at 0 h and from 37.2% to 61.6% at 24 h, while precision was maintained at 0 h (60.3% versus 59.6%) and decreased moderately at 24 h (43.4% versus 38.5%), so that the *F*_1_ score increased from 31.6% to 47.7% and from 40.1% to 47.4%, and the Jaccard index from 18.8% to 31.3% and from 25.1% to 31.1% (**Fig. 7C** and **Table S4**). The gain was largest at moderately edited sites: at sites edited at 10–20%, sensitivity increased from 13.0% to 38.6% (0 h) and from 22.9% to 54.4% (24 h; **Fig. 7D**). With the IVT control, in contrast, almost no Illumina site was recovered (precision 0.2% and 1.1%), because the G calls of the IVT RNA cancel the edited positions in the differential test. Thus, a large part of the inosine signal that the pileups miss is not absent but recorded as guanosine, and counting it substantially improves the detection of editing sites with an ADAR-deficient control.

## 3 Discussion

Nanopore basecallers now report, for every base of every read, a probability that it is modified. This probability is a model prediction learned from training data, and how well it identifies modified sites in a new biological sample has rarely been tested, and mostly for m6A and PsU ^29–31,47^. For A-to-I editing, our results show that the Dorado inosine model (sup@v6.0, SQK-RNA004) does not identify editing sites on its own. Useful results required a matched ADAR-deficient control, and even then Nanopore found at most about a third of the illumina reference sites that could be evaluated, while 40–57% of its calls were not in the Illumina reference. Nanopore sequencing can therefore contribute to the study of A-to-I editing, but only with an ADAR-deficient control, sufficient sequencing depth and, for now, confirmation by other methods.

The first problem is specificity. Confident inosine calls were as frequent in ADAR1-KO cells, which lack editing, as in WT cells, and even more frequent in IVT RNA, which contains no inosine (**Fig. 2**). The m6A model, tested on the same samples, performed better, but many of its confident calls were also found in the METTL3-KO control (**Fig. 2**). Editing is rare: only a few thousand of the millions of adenosines analysed are edited, most of them at low levels, so even a small error rate produces as many false calls as true ones. Similar problems have been reported for other Dorado modification models ^29–31^. The probability score assigned by the nanopore model alone therefore cannot separate edited from unedited positions, and editing sites have to be identified by comparison with a control.

The choice of this control is critical. IVT RNA, a common unmodified control, performed poorly: the number of sites it produced depended strongly on the probability threshold, and most of them were not supported by Illumina or by known editing catalogues (**Fig. 3**, **Fig. 4** and **Fig. S4**). Part of this may reflect the lower coverage of our IVT samples and features of IVT RNA that can mimic modifications ^48^. In addition, IVT RNA is not an exact unedited copy of the cellular RNA: since inosine is copied as G during its preparation ^12^, IVT RNA made from wild-type cells carries a G instead of an inosine at edited positions (**Fig. 7A**). The inosine model scores only bases that the basecaller recognise as A; at these reference adenosines, the reads carrying a G therefore receive no inosine score and are not counted in the IVT sample, which lowers its coverage precisely at the edited sites, in proportion to the editing level. For editing studies, an IVT control should therefore be made from ADAR1-KO RNA. A matched ADAR1-KO sample removes editing but keeps all the other modifications. Sites called against this control agreed with the Illumina reference twice as often as sites called against IVT RNA.

The second problem is sensitivity, and it has three causes. The first is depth: more than half of the missed reference sites in unstimulated cells and more than a third after stimulation had too few Nanopore reads to be evaluated (**Fig. 5C**), a limitation of the present dataset that deeper sequencing can solve. The second is the editing level: at most sites with enough reads, an inosine signal was present in wild-type but not in ADAR1-KO reads, yet in too few reads to be called (**Fig. 6**). Most editing sites are edited at low levels ^35,36^, and detection improved after stimulation, which raises editing (**Fig. 1A** and **Fig. S3**). The third is the basecaller: in about half of the edited reads the inosine was read as G and was therefore never scored by the inosine model, which scores only bases read as A (**Fig. 7**), most often in the sequence context preferred by ADAR (**Fig. S8A**). Counting these reads brought the Nanopore editing levels close to the Illumina levels after stimulation and nearly doubled the fraction of reference sites found with the ADAR1-KO control. With the IVT control it had the opposite effect: the true G of the IVT RNA was then counted as well, the IVT RNA looked as edited as wild-type RNA, and almost no site was found (**Fig. 7C**). Counting the G reads therefore helps only together with an ADAR-deficient control. Statistical methods that use per-read probabilities instead of fixed thresholds ^22^ could recover more sites.

Some Nanopore inosine calls were not in the Illumina reference. These are not necessarily errors, because the reference is not a ground truth. Some lay in repetitive regions and on unplaced genomic contigs, where long reads map better than short reads ^9,49^ (**Fig. S6**), and some were sites edited only after stimulation, which our reference excluded by design. Others showed a weaker ADAR sequence preference and may be false calls. Telling them apart will require methods that detect inosine chemically ^50^.

In practice, a Nanopore study of A-to-I editing needs an ADAR-deficient control from the same cells, enough depth at the sites of interest, and the inosine reads basecalled as G counted together with the inosine calls; even then, it will find mainly sites edited above roughly 10%. Novel sites, absent from short-read data and editing databases, should be confirmed independently. Models that also score inosine on reads called G, trained on data from ADAR-deficient systems, and standard analysis pipelines validated against independent data^30^ would address these problems. Our study has some limitations. The Illumina reference included only sites edited in both conditions, which favours specificity but leaves out sites edited only after stimulation; their effect is reported separately. The exact numbers will likely differ in other cell types, where editing levels differ. The requirements we identified (an ADAR-deficient control, sufficient depth, and counting inosine reads basecalled as G) are not specific to RAW 264.7 cells and should apply more generally. Finally, we evaluated the current Dorado inosine model; because these tools change quickly, new models should be evaluated in the same way, and our code and data are available for this purpose.

In conclusion, modification-aware basecalling makes inosine calls available for any direct RNA sequencing dataset, but these calls are not, by themselves, a measurement of A-to-I editing. With an ADAR-deficient control, enough depth and the reads read as G, Nanopore sequencing detects a reproducible subset of editing sites, including some in repetitive regions that short reads resolve poorly. Because it reads native molecules, it could in principle place modifications on individual transcripts, which becomes useful only once modification calls are reliable. For now, short-read sequencing remains the reference for identifying and quantifying A-to-I editing.

## 4 Methods

### Nanopore direct RNA sequencing data processing

Direct RNA sequencing reads were basecalled from the POD5 files and aligned with Dorado (v2.0.0) using the super-accuracy model sup@v6.0 together with the modification models for m5C, inosine, m6A, pseudouridine (*ψ*) and Nm of the four nucleotides (model names in the command below), discarding reads with a mean quality score below 6. Reads were aligned to the mouse reference genome (GRCm38/mm10) with the minimap2 aligner integrated in Dorado, in spliced mode without secondary alignments. Unmapped reads and alignments with a mapping quality below 10 were removed, and BAM files were sorted and indexed with SAMtools (v1.20) ^51^:

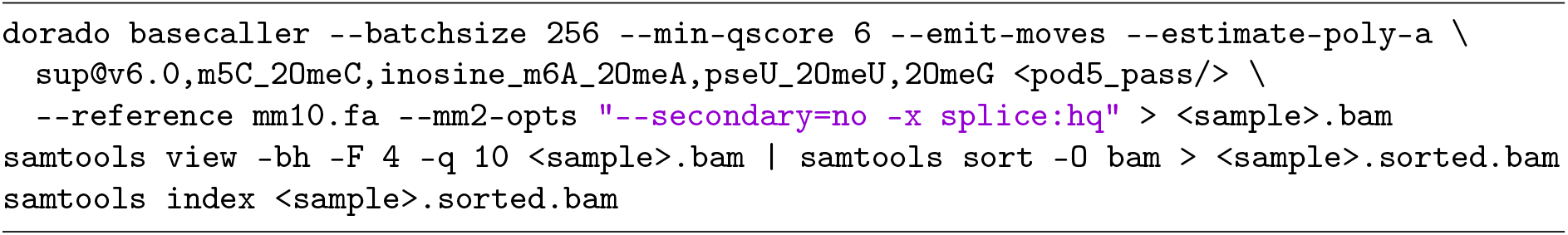

Per-site inosine calls were obtained with modkit (v0.6.4) pileup, restricted to adenosine and the inosine modification code (ChEBI 17596; `--modified-bases A:17596`), excluding the first and last 10 nucleotides of each read (`--edge-filter 10`) and using up to 20,000 reads per position (`--max-depth 20000 --high-depth`). The pass threshold for canonical adenosine calls was fixed at 0.7 (`--filter-threshold 0.7`), while the pass threshold for inosine calls (`--mod-threshold 17596`:*p*) was varied over *p ∈* {0.3, 0.5, 0.6, 0.7, 0.8, 0.9, 0.95, 0.99}, producing one pileup per sample and threshold. Pileups were generated for the 16 WT, ADAR1-KO and IVT samples, unstimulated (0 h) and after 24 h of IFN-*γ*/LPS stimulation:

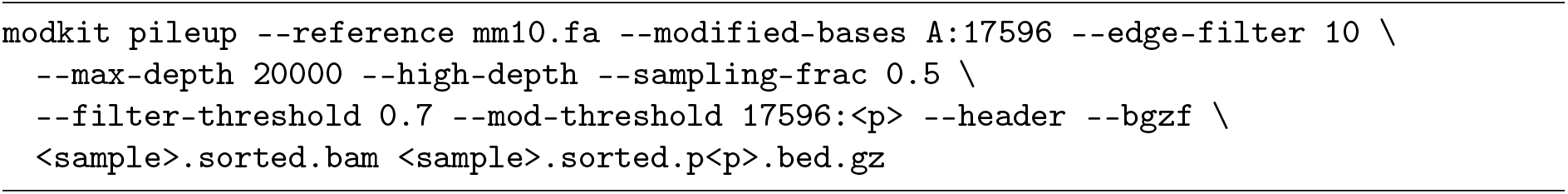

### Modification probability distributions

The specificity of the modification-aware models was assessed with modkit (v0.6.3) `modbam sample-probs` (`--hist, --sampling-frac 0.4`), which reports, for a random 40% of the reads of each sample, a histogram (bin width 1/256) of the probabilities of all base calls, for the 22 WT, ADAR1-KO, METTL3-KO and IVT samples:

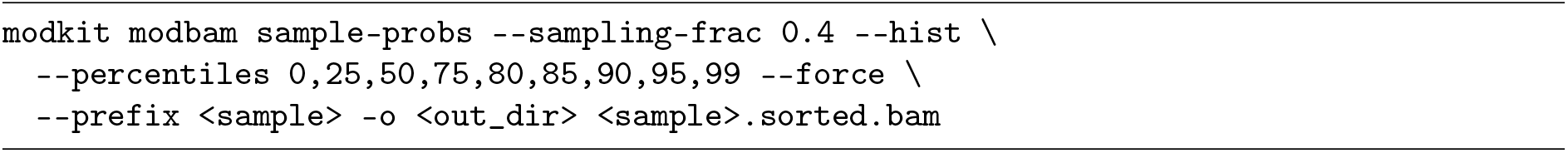

Each base call is assigned to the class with the highest probability (canonical base or one of the modifications called on that base). For m6A (code `a`), m5C (`m`) and inosine (`17596`), we counted, for each sample and probability cutoff, the calls of the modification with probability *≥* cutoff and divided them by all calls on the same base (canonical and modified):

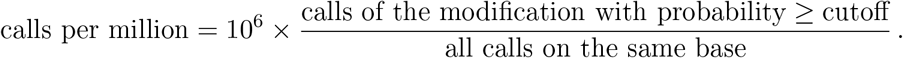

Values were averaged over the replicates of each group (WT, ADAR1-KO, METTL3-KO, IVT) and time point. Specificity was expressed as the enrichment in wild type relative to a negative control (METTL3-KO for m6A, ADAR1-KO for inosine, IVT for all three modifications) at the same cutoff and time point:

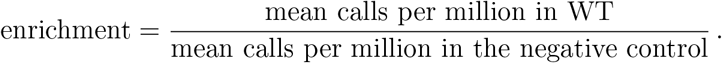

An enrichment close to 1 indicates that the negative control carries as many confident calls as wild type (**Fig. 2**; **Supplementary Data 1**).

### Differential inosine analysis (Nanopore)

Candidate inosine sites were identified in three steps (Steps 1–3), which were run independently for each of the eight pileup thresholds.

#### Step 1: per-sample positions

From each pileup, the inosine records (code 17596) with a valid coverage of at least 10 reads were extracted, keeping the position, the valid coverage and the number of modified reads. Four comparisons were defined, each contrasting a negative control with WT cells at the same time point: ADAR1-KO versus WT at 0 h and at 24 h, and IVT versus WT at 0 h and at 24 h. In the IVT-based comparisons, the IVT samples of both time points (2 + 2) were pooled as one group, as IVT RNA carries no modifications at either time point.

#### Step 2: filtering and statistical test

Within each comparison, a position was kept if (i) at least two samples carried at least one modified read, and (ii) it was covered (valid coverage *≥* 10) in at least two replicates of both groups. Differential modification was then tested per position with DSS^52^ (v2.54.0; `DMLtest`, `smoothing = FALSE`, as RNA modifications are discrete single-nucleotide events), using the number of modified reads and the valid coverage of each sample. P-values were corrected with the Benjamini–Hochberg procedure (FDR). The log_2_FC was computed from the group mean modification levels estimated by DSS as log_2_(*µ*_neg_) *−* log_2_(*µ*_WT_), so that negative values indicate a higher inosine level in WT than in the negative control. Positions with FDR *<* 0.05 and *|*log_2_FC*| ≥* 0.5 were considered significant in either direction; the group mean inosine levels of all tested positions are shown in **Fig. 3**, and the numbers of tested and significant positions per pileup threshold in **Fig. S2A–C** and **Supplementary Data 2**.

### Inosine site calling (Nanopore)

*Step 3: candidate sites.* Positions with FDR *<* 0.05 and log_2_FC *≤ −*0.5 at 0 h or at 24 h were pooled for each negative control (ADAR1-KO or IVT) and pileup threshold. At these positions, the inosine records of all samples of the comparison were extracted again from the pileups (WT and negative control, 0 h and 24 h), now with a valid coverage of at least 5 reads, and three filters were applied, in this order:

1. *Replicates:* the site was covered by at least 5 reads in at least two replicates of a condition, in at least two conditions, a condition being a sample group defined by genotype and time point (WT 0 h, WT 24 h, ADAR1-KO 0 h and ADAR1-KO 24 h, or IVT 0 h and IVT 24 h in the IVT-based comparison).
2. *Negative control:* the site was discarded if, in any negative-control condition (e.g. ADAR1-KO 0 h), the median valid coverage over replicates was *≥* 10 and the median modification frequency was *≥* 5%.
3. *Modified reads in wild type:* the site was retained only if, in at least one wild-type condition (WT 0 h or WT 24 h), the median valid coverage was *≥* 10 and the median number of modified reads was *≥* 5.

For the retained sites, valid coverage, modified-read counts and modification frequency were summarized as medians over the replicates of each condition; these values were used in all down-stream analyses. The number of positions removed by each filter is reported in **Supplementary Data 2**, and the number of candidate sites per pileup threshold in **Fig. S2D**.

### Illumina RNA-seq and A-to-I editing detection

Paired-end reads were trimmed with Trimmomatic (v0.38) ^53^, removing Illumina adapter sequences and discarding reads shorter than 50 nucleotides; only read pairs in which both mates survived trimming were kept. These were aligned to GRCm38/mm10 with STAR (v2.5.3a) ^54^, keeping uniquely mapped reads only and adding the XS strand attribute for spliced alignments:

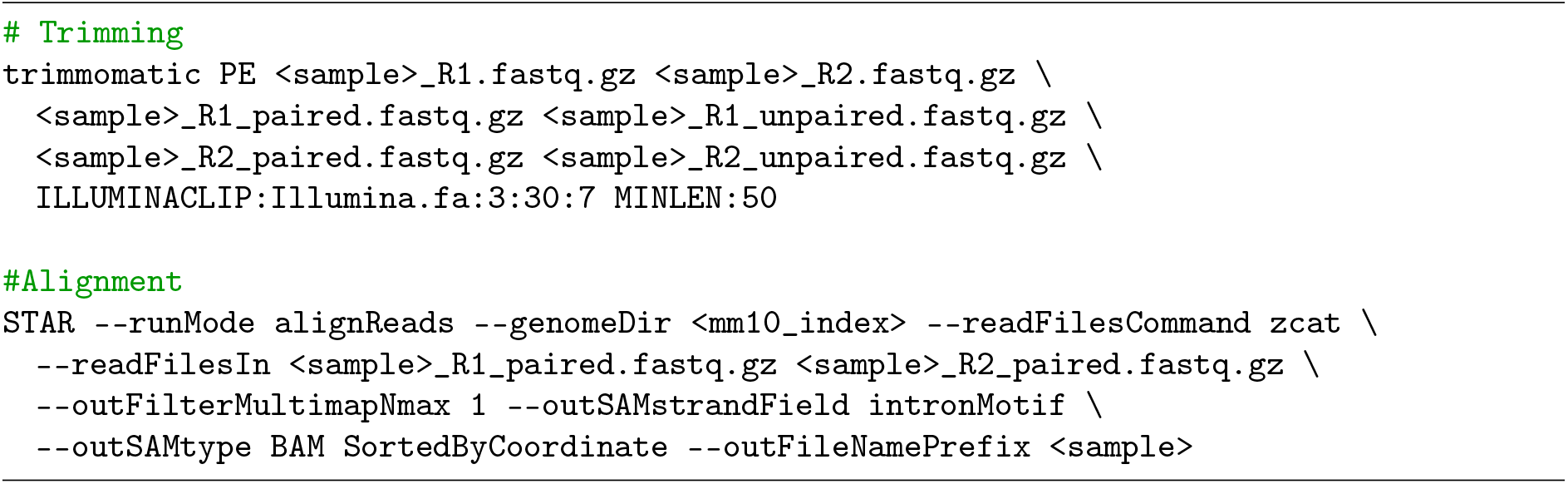

PCR duplicates were removed with SAMtools (`fixmate -m, markdup -r -s`). RNA editing candidates were called on the deduplicated alignments with REDItools (v2.0)^55,56^ in strandaware mode (`-s 2`, strand confidence mode 2, `-T 2`), with a minimum mapping quality of 25 (`-q 25`) and a minimum base quality of 35 (`-bq 35`), excluding the first and last 10 nucleotides of each read (`-mbp 10 -Mbp 10`) to avoid artefacts at read ends, and masking homopolymeric regions of at least 4 nucleotides (`-os 4`) and positions near splice sites (mm10 splice-site annotation, `-sf`):

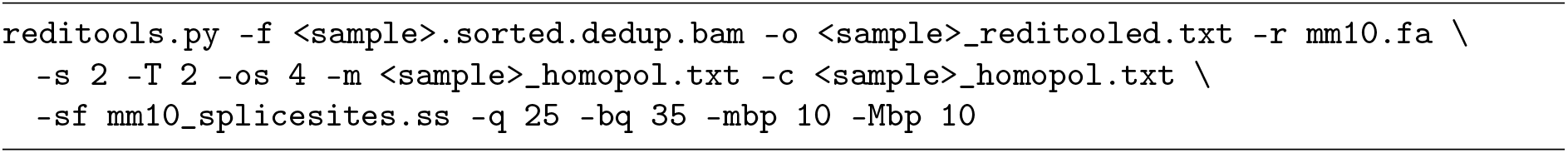

The REDItools output was restricted to positions with reference base A on the transcribed strand (REDItools reports the reference base and base counts on the inferred strand).

To remove positions that differ from the reference for reasons other than A-to-I editing (genomic variants of the cell line, spontaneous adenosine deamination, sequencing or alignment errors), a background of non-variant positions was built from ADAR1-knockout samples (0 h and 24 h): positions with no substitution and a coverage (base quality *≥* 30) of at least 10 reads, detected in at least two of three replicates of both time points. Background positions with a median coverage below 10 were removed. A-to-G substitutions (A*→*G on the transcribed strand) with a coverage of at least 5 reads and a single substitution type were then called in the wild-type samples (0 h and 24 h) at these background positions. Editing sites were kept if they were detected in at least two of three replicates at both time points, and if the median coverage over replicates was *≥* 10 and the median number of edited (G) reads was *≥* 5 in at least one time point (2,971 editing sites; **Fig. 1**). Requiring detection at both time points favours specificity but excludes editing sites that are specific to stimulated cells. Positions with A-to-G substitutions at 24 h only were rarely listed in REDIportal (7.6%, compared with 46.7% of the retained sites) and rest on two or three replicates of a single time point; they were analysed separately (see *Stimulation-specific editing sites*).

### Stimulation-specific editing sites

To quantify this effect, the excluded positions were reconstructed with the same criteria, except that detection at one time point was sufficient: A-to-G substitutions at the background positions in at least two of three wild-type replicates at 24 h but not at 0 h, with a median coverage *≥* 10 and a median of *≥* 5 edited reads at 24 h, not contained in the reference set (5,231 positions, median editing frequency 5%, 7.6% in REDIportal; **Supplementary Data 2**). These positions were compared with the Nanopore candidate sites at the identical position. In a sensitivity analysis, the 24 h benchmark (see below; pileup threshold 0.8, coverage *≥* 10, frequency *≥* 5%, window 0) was repeated with three references: the original Illumina sites, the original sites extended by the 24 h-only positions, and the 24 h-only positions alone (in the last case, only Nanopore calls not contained in the original reference were scored). Read depth at the 24 h-only positions was measured as for all benchmarked positions.

### Read depth at the benchmark sites

To evaluate whether a site could be detected by each platform, read depth was measured at all benchmarked positions: REDIportal sites, Illumina editing sites and Nanopore inosine calls of every comparison and pileup threshold, and the 24 h-only Illumina positions (137,594 positions, extended by *±*2 nt to allow window-based matching and merged into one BED file). Depth was computed per sample with SAMtools (v1.20) `depth` on the deduplicated Illumina BAM files of the WT and ADAR1-KO samples (the input of REDItools) and on the filtered Nanopore BAM files of the WT, ADAR1-KO and IVT samples, at 0 h and 24 h, reporting also positions without reads (-a), so that zero depth could be distinguished from unmeasured positions, and counting only reads with a mapping quality of at least 25 (-Q 25):

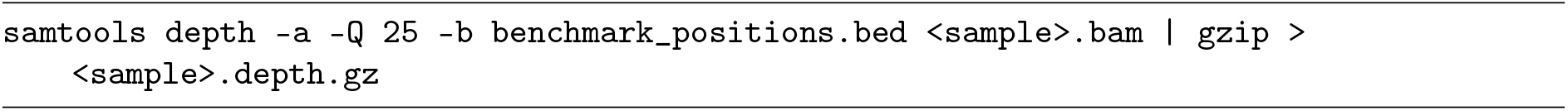

For each platform, sample group and condition, the median depth over replicates was used. Depth obtained with SAMtools correlated with the REDItools coverage at Illumina editing sites (Spearman *ρ* = 0.997; **Supplementary Data 2**).

Depth was compared between platforms using the fraction of sites with depth above a given value per sample, the Spearman correlation of depth between all samples, and the per-site Illumina versus Nanopore depth (Spearman *ρ* and median Nanopore/Illumina ratio). Differences in depth between the platforms were expressed per site as

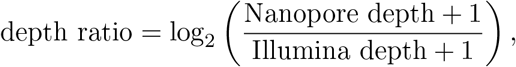

which is 0 for equal depth and negative where Nanopore has fewer reads. The depth ratio was plotted against the mean depth of the two platforms (mean of log_10_(depth + 1); Bland–Altman plot ^57^), with a smoothed trend fitted by a generalised additive model (cubic regression spline; R package mgcv), and its median was computed per chromosome over the sites covered by both platforms (**Fig. S7**).

To relate depth to detection, editing sites were classified by identical position as detected by both platforms, by Illumina only or by Nanopore only (ADAR1-KO-based candidate sites, pileup threshold 0.8), and the wild-type depth of both platforms was compared between these classes (**Fig. 5A–B**). Illumina editing sites of each condition (at least two replicates with coverage *≥* 10 and editing frequency *≥* 5%) that were not called by Nanopore were assigned to the first applicable cause: Nanopore wild-type depth *<* 10, Nanopore ADAR1-KO depth *<* 10, or sufficient depth in both but no Nanopore call (**Fig. 5C**). For the last group, the Nanopore wild-type depth and the Illumina editing frequency (median over replicates) were compared with those of the Illumina sites detected by Nanopore, reported as medians and the difference between the medians. Because neighbouring sites in the same transcript are not independent observations, these comparisons are descriptive and no p-values are given. For both groups, the expected number of edited Nanopore reads was computed as

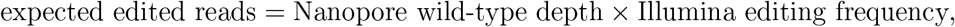

which estimates how many Nanopore reads would carry inosine at the site if both platforms measured the same editing level, and summarized as the fraction of sites with fewer than 5 expected edited reads (the minimum number of modified reads required for a call) (**Fig. 5D–E**; **Supplementary Data 2**).

### Overlap between Nanopore and Illumina editing sites

The Nanopore candidate sites of each comparison (pileup threshold 0.8), the Illumina editing sites and the REDIportal sites^44,45^ (mm10, version 3) were compared at the identical genomic position (0-based start coordinate), without coverage restriction. For each Nanopore site set we report the number and fraction of sites that coincide with an Illumina editing site or a REDIportal site; for the Illumina sites, the fraction contained in REDIportal (**Fig. 4A–B**).

### Per-read inosine probabilities at detected and missed sites

Illumina editing sites that were detected by Nanopore, or covered by at least 10 reads in both the Nanopore wild-type and ADAR1-KO samples but not called (ADAR1-KO-based sites, pileup threshold 0.8), were extended by *±*5 nt on the edited strand (1,930 windows). Per-read modification probabilities in these windows were extracted from the Dorado BAM files of the WT, ADAR1-KO and IVT samples (0 h and 24 h) with modkit (v0.6.4) `extract full`, restricted to these windows and to mapped reads and excluding the first and last 10 nucleotides of each read, as in the pileups:

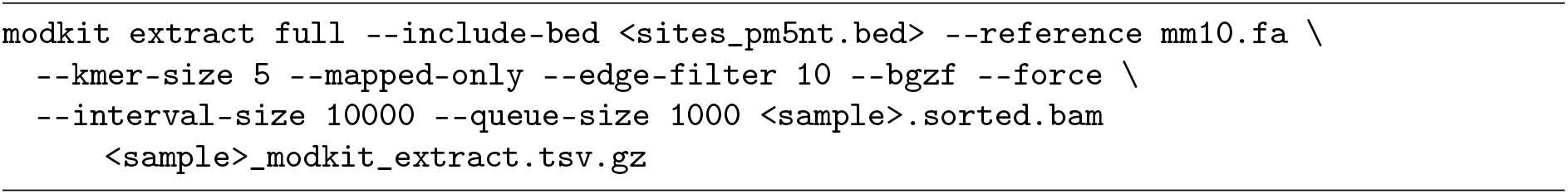

`extract full` reports the modification probabilities of every read and position without applying any threshold; positions without an explicit probability (implicit canonical calls in the MM tag) were kept as canonical. To make the per-read results comparable with the site calls, the pileup thresholds were then applied in R to every read and adenosine: the canonical probability was computed as 1 *− P* (m^6^A) *− P* (I) *− P* (Am), the most likely class was called if its probability was *≥* 0.8 (inosine) or *≥* 0.7 (all other classes), and positions below these thresholds were discarded. The distribution of the per-read inosine probability at the edited adenosine (**Fig. 6D**) was computed from the unfiltered probabilities. Reads of the replicates were pooled per genotype and time point, and sites with at least 10 passing reads were analysed. The fraction of inosine calls at the edited adenosine was compared between WT and ADAR1-KO at the same sites as the median of the per-site difference and the fraction of sites with a higher level in WT (descriptive, as above) (**Fig. 6A**). The difference in inosine call fraction between WT and ADAR1-KO was also computed at each position of the *±*5 nt window and summarized as mean and 95% confidence interval (*±*1.96 standard errors) over sites (**Fig. 6B**). The WT inosine call fraction was compared with the Illumina editing frequency by Spearman correlation (**Fig. 6C**).

### Inosine reads basecalled as guanosine

The Dorado modification model scores inosine only on bases called A; reads in which the base-caller called G at a reference adenosine therefore carry no inosine probability, and modkit pileup counts them neither as modified nor in the valid coverage (they are included in `count_diff`, together with A-to-C and A-to-U calls). These reads were counted per position and strand with SAMtools (v1.20) mpileup, using primary alignments only and no base-quality, mapping-quality or depth filter, one chromosome at a time:

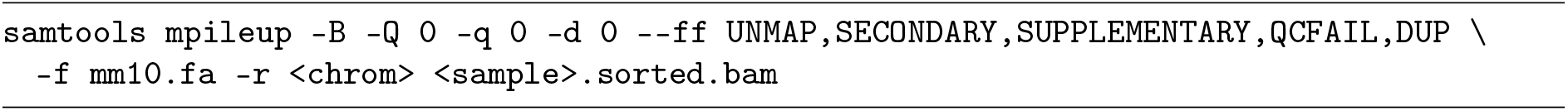

Because direct RNA reads are aligned in the orientation of the transcript, reads basecalled G at a reference A were counted on forward-strand alignments (+ strand), and reads basecalled C at a reference T on reverse-strand alignments (*−* strand). Unlike the pileups, mpileup does not exclude the 10 nucleotides at the read ends (0.2% more G reads in a 10-Mb test region). In a test region, the counts were identical to those obtained from the per-read output of modkit `extract full`.

To count these reads as modified, the reads basecalled G were added both to the modified reads and to the valid coverage of the inosine rows of each pileup (threshold 0.8), and the modification frequency was recomputed:

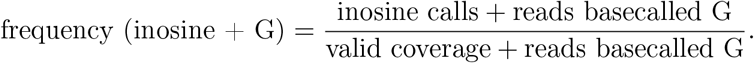

Only positions present in the pileup were changed. The differential analysis, the site calling and the benchmarking against Illumina were then repeated unchanged with these pileups (*Differential inosine analysis*, *Inosine site calling*, *Benchmarking*). The read depth (*Read depth at the benchmark sites*) was measured again with the new candidate sites added to the position list; the depth at all other positions, and thus all other benchmark results, were unchanged.

At the 2,971 Illumina editing sites and at 10,000 randomly chosen adenosines (valid coverage *≥* 20 in the first WT 24 h replicate, Illumina sites excluded), the reads called A with a passing call, the inosine calls and the reads basecalled G were pooled over the replicates of each genotype and time point. For Illumina sites with an unknown strand, the strand with more WT reads was used. The percentage of reads basecalled G (reads basecalled G divided by all reads called A or G) was computed per replicate over all sites with *≥* 10 reads (**Fig. 7A**). The fraction of the edited reads basecalled G was computed from the excess of WT over ADAR1-KO reads, pooled over the sites of a group (all sites, or sites sharing the 5*^′^* and 3*^′^* neighbour on the RNA strand; sites with *≥* 10 reads in both genotypes). In each genotype, the fraction of reads basecalled G and the fraction of reads with an inosine call were computed over all reads called A or G, and the excess of WT over ADAR1-KO was taken as the edited reads:

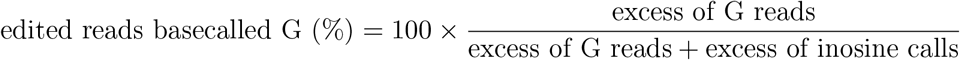

(**Fig. S8A**). The WT editing level was computed either from the inosine calls only (inosine calls divided by the reads called A) or with the reads basecalled G, as in the formula above, and compared with the Illumina editing frequency (median over replicates; sites detected in at least two replicates) by Spearman correlation and the slope of a linear fit through the origin (**Fig. 7B**). Because these measure association rather than agreement, the same sites were also compared with Lin’s concordance correlation coefficient^46^

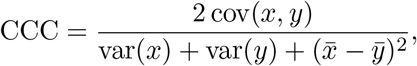

with *x* the Illumina editing frequency and *y* the Nanopore level, which equals 1 only if all sites lie on the identity line (computed with variances over *n* as in the original definition, using the R package yardstick; 95% confidence interval from 1,000 bootstrap resamples of sites), and with a Bland–Altman analysis^57^ using Illumina as reference: the difference between the Nanopore and the Illumina level was plotted against the Illumina level, and its mean (bias) and mean *±*1.96 standard deviations (limits of agreement) were reported (**Fig. S8C**). These analyses are descriptive. For **Fig. S8B**, the per-site WT minus ADAR1-KO level was averaged within Illumina frequency classes and expressed as a percentage of the mean Illumina frequency of the same sites. Precision, sensitivity and *F*_1_ score were computed as described below (coverage *≥* 10, frequency *≥* 5%, identical position; **Fig. 7C**); sensitivity was additionally computed within Illumina frequency classes (**Fig. 7D**). Results with inosine calls only reproduced the main benchmark exactly.

### Annotation and characterisation of editing sites

For the Illumina editing sites, editing frequency and coverage were summarized as medians over the replicates of each sample group. The editing frequency in WT and ADAR1-KO cells was compared per site at 0 h and at 24 h (**Fig. S1B–C**). The distribution of the WT editing frequency before and after stimulation (**Fig. 1A**) was computed for the sites with a median coverage *≥* 10 and *≥* 5 edited reads, detected in at least two replicates at both time points; the corresponding distribution of the Nanopore WT inosine frequency at the ADAR1-KO-based candidate sites is shown in **Fig. S3A**. The sequence context of the editing sites (*±*5 nt on the edited strand reported by REDItools, extracted from mm10 with BSgenome) was summarized as base frequencies minus 0.25 per position (**Fig. 1B**).

Illumina editing sites and Nanopore candidate sites were annotated with HOMER ^58^ (mm10), using 0-based BED coordinates; transposable elements (SINE, LINE and LTR) and repeat families were taken from the repeat annotation of the HOMER mm10 package (v7.0), which is derived from the UCSC RepeatMasker track. Sites on unplaced genomic contigs, which HOMER does not annotate, were reported separately (**Fig. 1C**). Sites detected by both platforms, by Nanopore only or by Illumina only (identical position; ADAR1-KO-based Nanopore sites, pileup threshold 0.8) were compared by genomic region, repeat family and sequence context (*±*5 nt on the edited strand, extracted from mm10 with BSgenome; base frequencies minus 0.25; **Fig. S6**). Overlap with known polymorphisms was tested at the identical position against the Mouse Genomes Project v5 SNP calls annotated with dbSNP build 142 (36 strains); a site was considered a SNP of the RAW 264.7 background if BALB/cJ carried the alternative allele (genotype passing the strain-specific filters; **Fig. S1A**). Editing frequencies before and after stimulation (medians over wild-type replicates) were compared between platforms at the sites detected by both, requiring at least 10 reads in both platforms at the respective time point (Spearman correlation; **Fig. S3B**).

### Benchmarking of Nanopore inosine calls against Illumina

Each Nanopore call set (comparison *×* pileup threshold) was compared with the Illumina editing sites of the same condition (0 h or 24 h). For both platforms, wild-type sites were summarized per condition as medians over replicates and filtered with the same criteria: modification/editing frequency *≥* 5% and coverage *≥ X*, with *X ∈ {*10, 20, 30, 40, 50, 60*}*. Illumina sites additionally had to be detected in at least two replicates. To avoid counting sites that a platform could not observe, an Illumina site was used as a true site only if the Nanopore wild-type depth was *≥ X* and the depth of the corresponding Nanopore negative control (ADAR1-KO of the same condition, or IVT pooled over both conditions) was *≥* 10; likewise, a Nanopore call was scored only if the Illumina wild-type depth at that position was *≥ X*. Nanopore calls were matched to Illumina sites on the same chromosome and a compatible strand at the identical position (window 0 nt; windows of up to *±*2 nt were evaluated in a sensitivity analysis). Matching was one-to-one: each call could be assigned to at most one site and vice versa, assigning the closest pairs first. True positives (TP; reference sites called by Nanopore) were matched pairs, false positives (FP; Nanopore calls not in the reference) unmatched Nanopore calls and false negatives (FN; reference sites not called by Nanopore) unmatched Illumina sites, so that TP + FP equals the number of calls and TP + FN the number of reference sites. Performance was summarized as

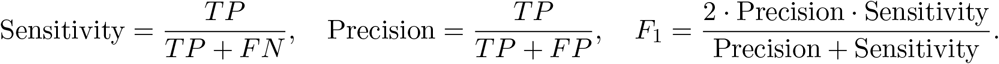

True negatives were not defined, as no set of verified non-edited positions is available. Because the Illumina sites are a reference and not a ground truth, FP and FN denote Nanopore calls outside the reference and reference sites missed by Nanopore rather than verified errors. We therefore also report the Jaccard index, *TP/*(*TP* + *FP* + *FN*), the fraction of all sites detected by either platform that are detected by both, which treats the two platforms symmetrically.

Metrics at coverage *≥* 10 are shown in **Fig. 4C** and **Table S4**, and for all coverage cut-offs in **Fig. S4A–B** (**Supplementary Data 2**). Because each negative control required only its own depth, the ADAR1-KO- and IVT-based calls were scored on different sets of Illumina sites. To compare the two controls on the same sites, the benchmark was repeated with Illumina sites and Nanopore calls both restricted to positions with a depth *≥* 10 in the Illumina wild-type samples and in the Nanopore WT, ADAR1-KO (same condition) and IVT (pooled) samples, with all other rules unchanged (**Fig. S5**; **Supplementary Data 2**). The differential analysis, site calling and benchmarking were performed independently for each of the eight pileup thresholds; results are reported in the main text for threshold 0.8, which gave the highest *F*_1_ score for the ADAR1-KO-based calls after stimulation and lay within a broad *F*_1_ plateau (0.7–0.9), and for all thresholds in the Supplementary Information.

### Benchmarking against REDIportal

Illumina editing sites and Nanopore calls were additionally compared with the mouse A-to-I editing sites of REDIportal^44,45^ (mm10, version 3; 107,094 sites). As this catalogue provides no tissue or cell-type information and is not specific to macrophages, the analysis was restricted to positions covered by at least 10 reads (median over replicates) in the wild-type samples of both platforms in the respective condition; both the REDIportal^44,45^ sites (true sites) and the calls of each platform were limited to these positions, without the negative-control depth requirement of the Illumina benchmark. Calls were filtered as in the Illumina benchmark (frequency *≥* 5%, coverage *≥* 10, at least two replicates for Illumina) and matched one-to-one at the identical position on a compatible strand. Metrics were computed as above, using Nanopore pileup threshold 0.8 for the comparison between platforms (all thresholds in **Fig. S4C**; **Supplementary Data 2**).

### Data Generation

All sequencing data used in this study were generated as part of a larger project. Detailed information regarding cell culture conditions, treatments, and sequencing procedures can be found in Griesche, V. & Di Giorgio, S. et al ^37^. The samples analysed in this study and their numbers of sequenced and aligned reads are listed in **Table S1**.

## Data availability

For mouse mRNA analyses, we used publicly available datasets from the NCBI BioProject database with project ID PRJNA1232413 (Illumina RNA-seq, and Nanopore direct RNA sequencing kit: SQK-RNA004, PromethION flow cell: FLO-PRO004RA) ^37^. Processed data are available at EMBL-EBI BioStudies under accession S-BSST3486: REDItools tables of the wildtype and ADAR1-KO samples (0 h and 24 h), modkit pileups of all Nanopore samples at the eight inosine pass thresholds, the Illumina editing sites, the Nanopore candidate inosine sites and the read depth at the benchmarked positions. The mouse A-to-I editing sites of REDIportal (mm10, version 3) were obtained from the REDIportal database. The code used for the analyses is available at https://github.com/Salvobioinfo/AtoI-nanopore-vs-illumina.

## Software

Analyses were performed in R (v4.4.3) with data.table (v1.17.0), dplyr (v1.2.1), DSS (v2.54.0), bsseq (v1.42.0), GenomicRanges (v1.58.0), Biostrings (v2.74.1), BSgenome (v1.74.0, with the `BSgenome.Mmusculus.UCSC.mm10` package), VariantAnnotation (v1.52.0), biomaRt (v2.62.1), ggplot2 (v4.0.1), ggseqlogo (v0.2), mgcv (v1.9.1), patchwork (v1.3.0) and yardstick (v1.4.0; concordance correlation coefficient), and with Dorado (v2.0.0), modkit (v0.6.4 for `pileup` and `extract`, v0.6.3 for `sample-probs`), SAMtools (v1.20), STAR (v2.5.3a), Trimmomatic (v0.38), REDItools (v2.0) and HOMER (v5.1, mm10 package v7.0).

## Acknowledgements

We thank the Next Generation Sequencing Core Facility of the German Cancer Research Center (DKFZ) for providing excellent sequencing services, and the Omics IT and Data Management Core Facility (ODCF) of the DKFZ for computing infrastructure and support. We thank Prof. Nina Papavasiliou, Dr. Riccardo Pecori, Dr. Silvestro Conticello and Prof. Julia Brettschneider for their feedback on the manuscript. We gratefully acknowledge funding by the TRR319 RMaP (V.G. and S.D.G.). S.D.G. is financed by the Baden-Württemberg Stiftung gGmbH, project BWST-ISF2025-050, EpiCode-Macro

## Author Contributions

S.D.G. conceived the study. V.G. generated the clonal knockout RAW 264.7 cell lines and the IVT RNA and performed the other experiments. S.D.G. pre-processed the Nanopore sequencing data and performed all bioinformatic analyses of the Illumina and Nanopore sequencing data. S.D.G. provided scientific guidance. S.D.G. wrote the manuscript.

## Computational tools and AI-assisted methods

R (v4.4.3; packages listed above) was used for data analysis, statistical analysis and data visualization. Artificial intelligence–assisted tools were used to support code development, language refinement and proofreading of the manuscript. All scientific content, analyses, interpretations and conclusions remain the responsibility of the authors.

## Acronyms

ADAR1-KO: ADAR1 knockout. 1–13, 15–20, 22, 23, 25–29
BAM: Binary Alignment Map. 12, 22, 25, 26
ChEBI: Chemical Entities of Biological Interest. 22
dRNA-seq: direct RNA sequencing. 2, 22
G: guanosine. 3
I: inosine. 2, 5, 6, 8, 9, 11–13, 16–23, 25–29
IVT: *in vitro* transcribed. 1, 2, 5, 6, 8–10, 12, 13, 16–20, 22, 23, 25, 26, 28–30
log_2_FC: log2 fold change. 23
m1A: N1-methyladenosine. 2
m5C: 5-methylcytosine. 2, 5, 22
m6A: N6-methyladenosine. 2, 5, 20, 22
METTL3-KO: METTL3 knockout. 5, 12, 20, 22
Nm: 2*^′^*-O-methylation. 2, 22
ONT: Oxford Nanopore Technology. 2, 10
PsU: pseudouridine (*ψ*). 2, 20, 22
SNPs: single nucleotide polymorphisms. 3
WT: wild-type. 1–6, 8–13, 15–20, 22, 23, 25–29

## 5 Supplementary Information

**Figure S1:**
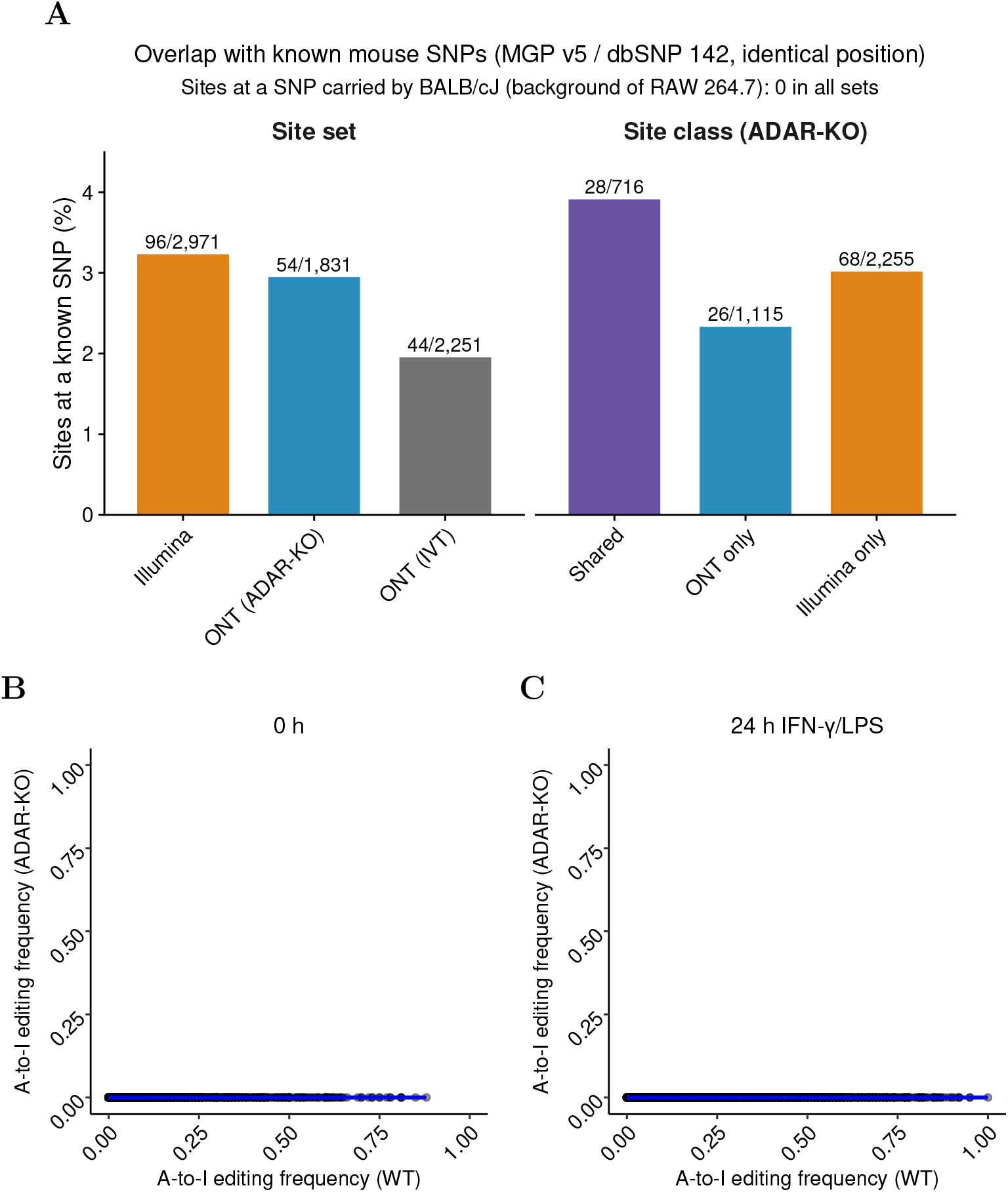
Overlap of editing sites with known mouse SNPs and editing frequency in ADAR1-KO cells. **(A)** Percentage of sites located at the position of a single-nucleotide polymorphism of the Mouse Genomes Project (v5, 36 strains, dbSNP build 142), for the Illumina editing sites, the Nanopore candidate sites obtained with the ADAR1-KO or IVT control (pileup threshold 0.8; left), and the sites detected by both platforms or by only one of them (ADAR1-KO-based; right). Numbers give sites at a SNP over all sites of the set. No site coincided with a SNP carried by BALB/cJ, the genetic background of RAW 264.7. **(B, C)** A-to-I editing frequency of each of the 2,971 Illumina editing sites (Fig. 1) in WT (x axis) and ADAR1-KO cells (y axis) in unstimulated cells (0 h; B) and after 24 h of IFN-*γ*/LPS stimulation (C); medians over three replicates. ADAR1-KO frequencies are 0 by construction of the site selection. Blue line, linear fit. Values of A are given in Supplementary Data 2, the editing sites (B, C) in Supplementary Data 1. See Methods, *Annotation and characterisation of editing sites*.

**Figure S2:**
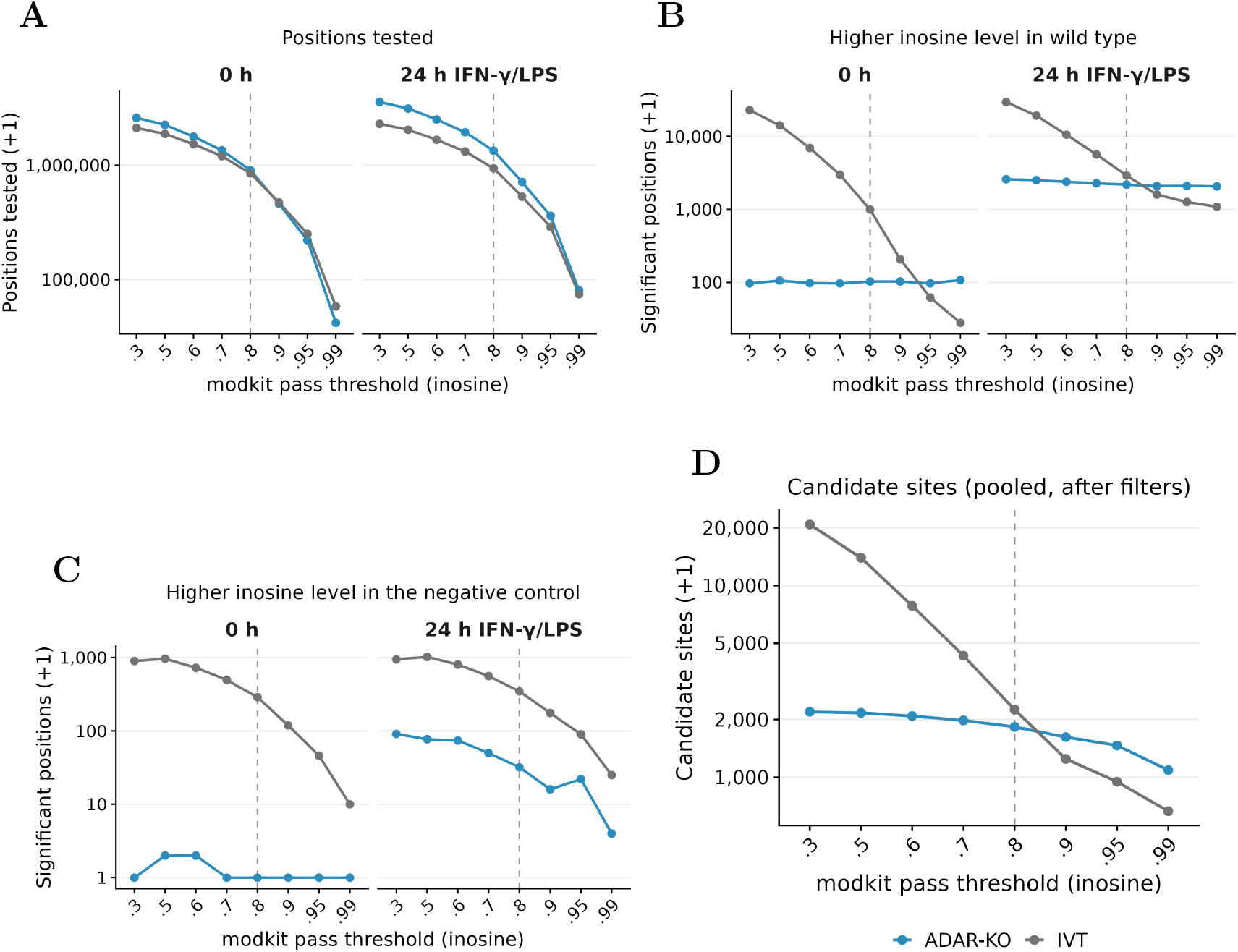
Dependence of the differential inosine analysis on the modkit pass threshold. The differential analysis (DSS) between wild-type cells and the ADAR-KO (blue) or IVT (grey) negative control was repeated for eight modkit pass thresholds of inosine calls (0.3–0.99). **(A)** Positions tested. **(B)** Positions with significantly higher inosine levels in wild type (FDR *<* 0.05, log_2_FC *≤ −*0.5). **(C)** Positions with significantly higher inosine levels in the negative control (FDR *<* 0.05, log_2_FC *≥* 0.5), which cannot reflect editing. **(D)** Candidate editing sites after pooling both time points and filtering. (A–C) are shown separately for unstimulated cells (0 h) and after 24 h of IFN-*γ*/LPS stimulation. y axes are on a log_10_ scale (counts +1; gridlines at powers of ten, so that, for example, the highest point in B, 22,778 positions for IVT at 0 h and threshold 0.3, lies above the 10,000 line); the dashed line marks the threshold used in the main figures (0.8). Values are given in Supplementary Data 2. See Methods, *Differential inosine analysis* (A–C) and *Inosine site calling* (D).

**Figure S3:**
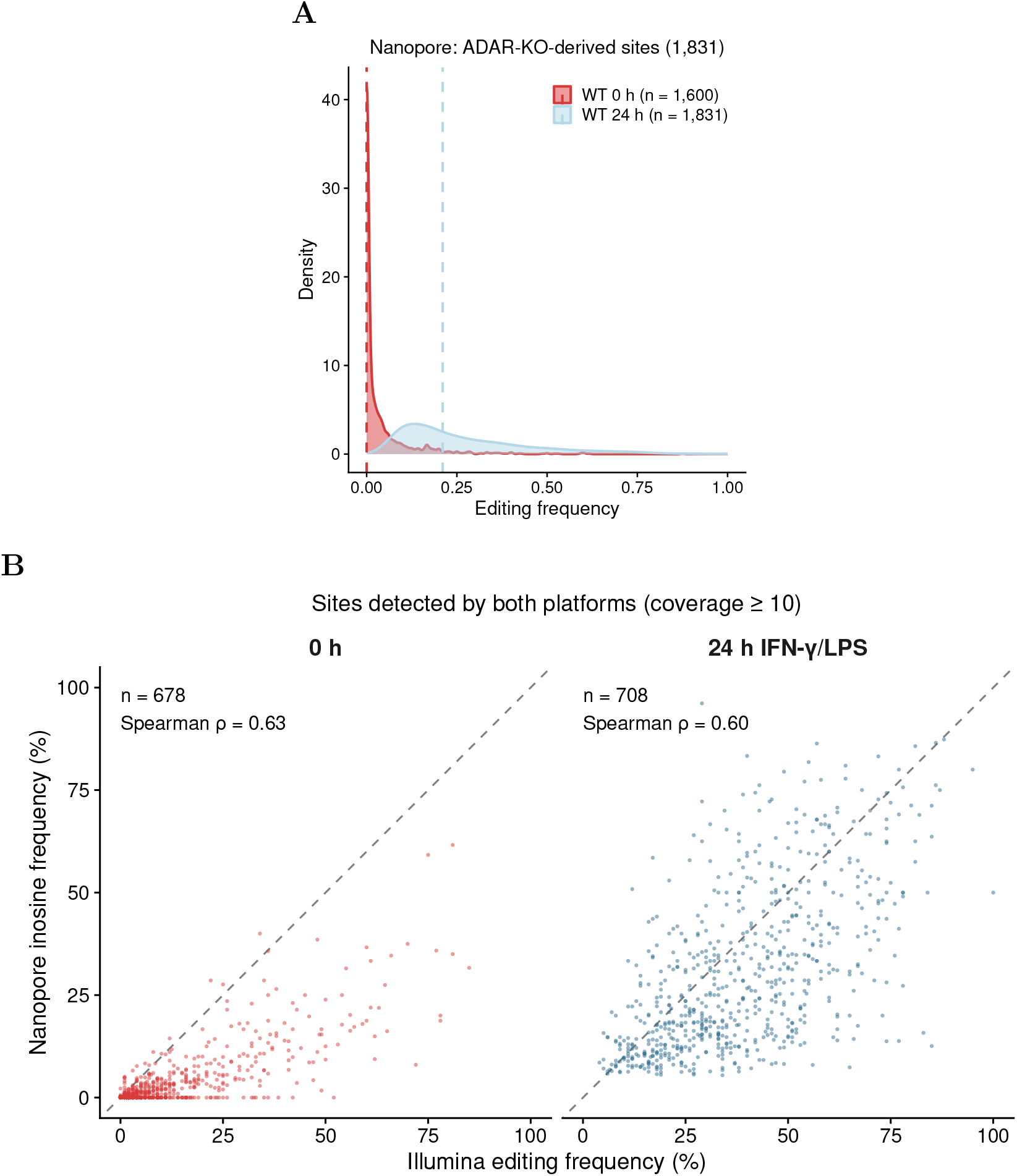
Stimulation-induced editing measured by both platforms. **(A)** Distribution of the inosine frequency (Nanopore, modkit pass threshold 0.8) at the 1,831 candidate sites obtained with the ADAR1-KO control in WT cells before (0 h, red) and after 24 h of IFN-*γ*/LPS stimulation (blue).**(B)** Editing frequency measured by Nanopore versus Illumina at the sites detected by both platforms, in unstimulated cells (0 h; 678 sites) and after 24 h of stimulation (708 sites); sites with *≥* 10 reads in both platforms at the respective time point. Dashed line, identity; *ρ*, Spearman correlation. Values are given in Supplementary Data 2. See Methods, *Annotation and characterisation of editing sites*.

**Figure S4:**
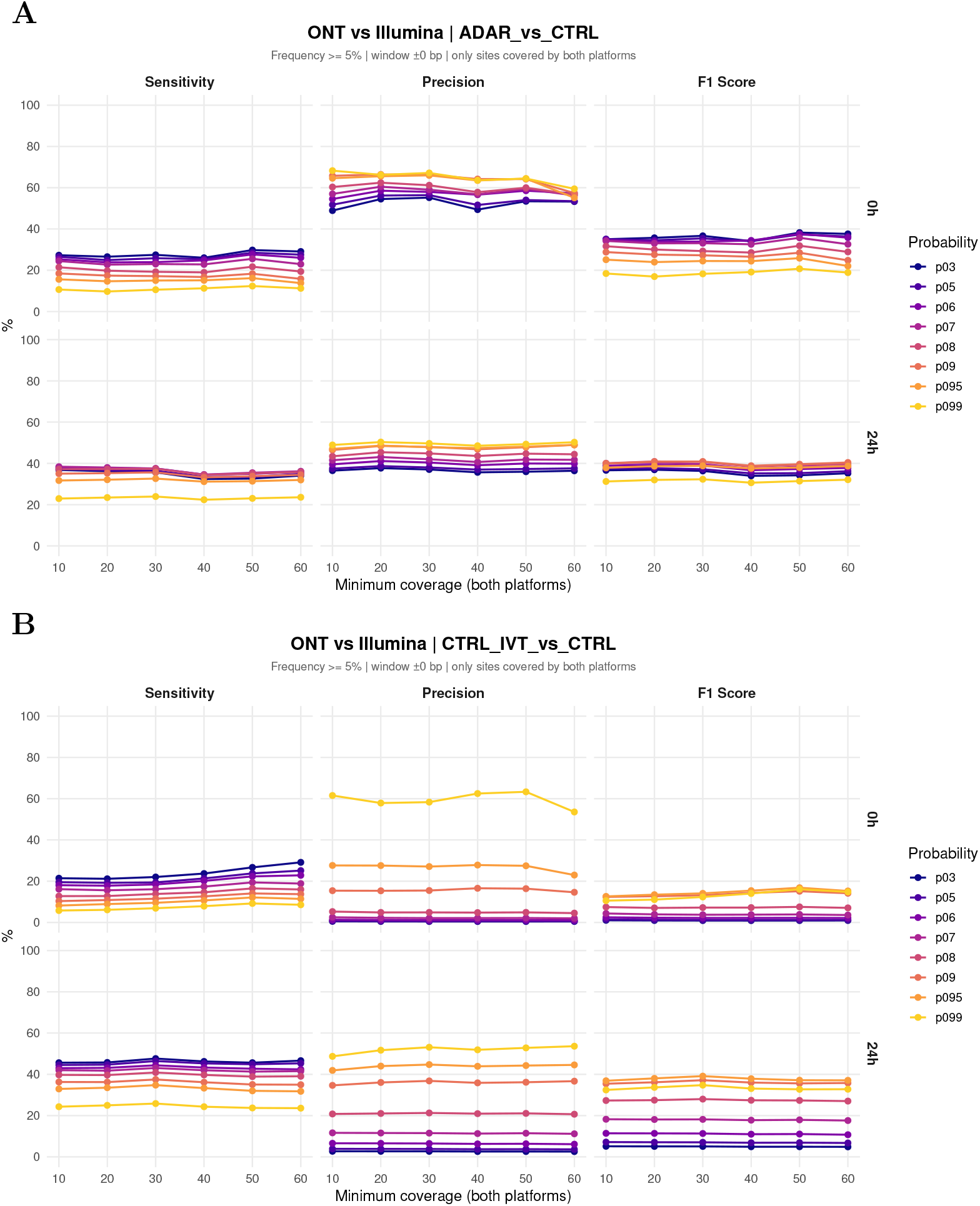
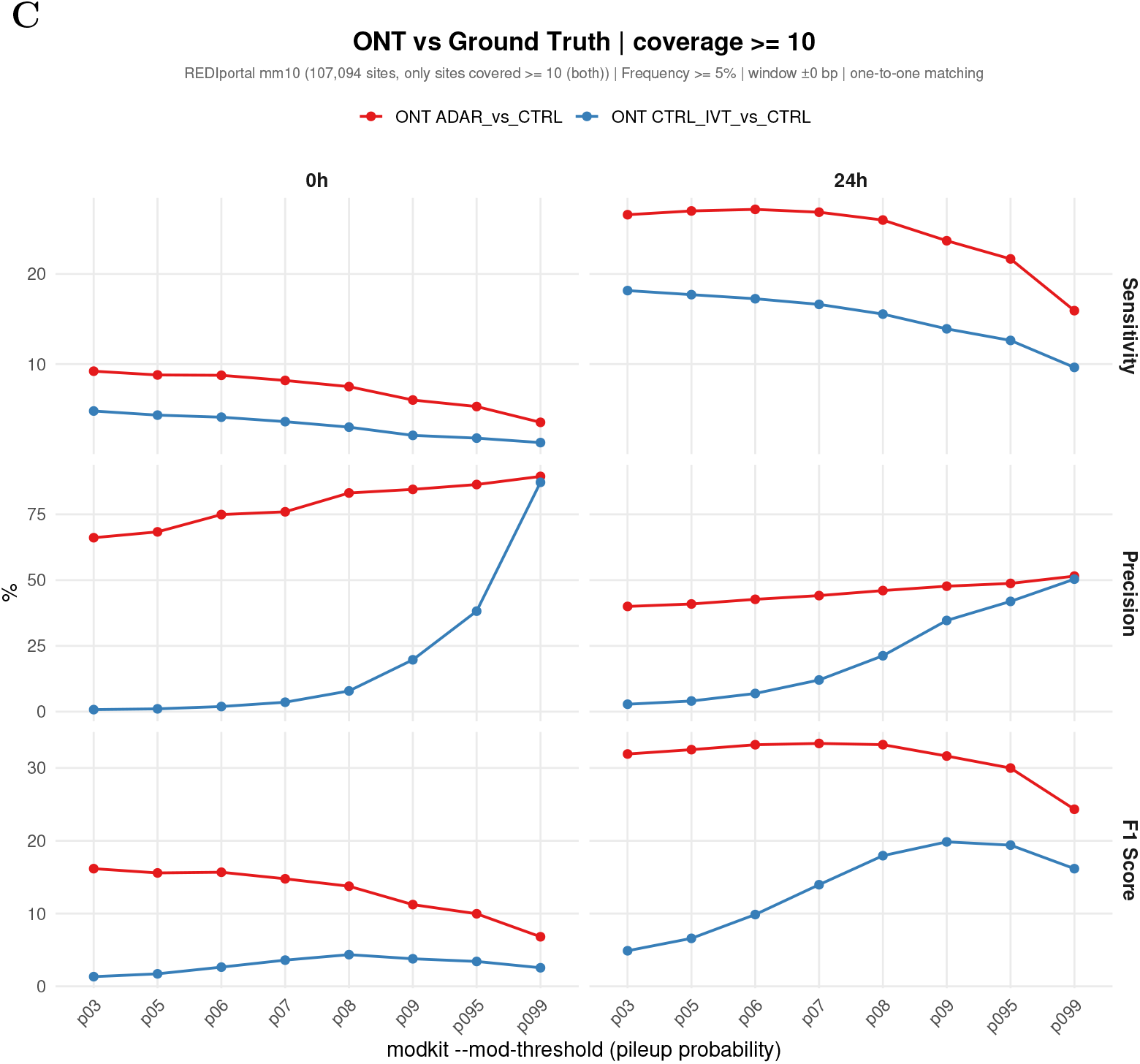
Robustness of the Nanopore–Illumina benchmark. **(A, B)** Precision, sensitivity and *F*_1_ score of Nanopore candidate sites obtained with the ADAR1-KO (A) or IVT (B) negative control, relative to the Illumina editing sites, as a function of the minimum read depth required in both datasets (10–60 reads), for each modkit pass threshold. Only sites covered in both datasets and in the Nanopore negative control were evaluated (editing frequency *≥* 5%, identical position, one-to-one matching). **(C)** Precision and sensitivity of the Nanopore candidate sites relative to the REDIportal catalogue of known mouse editing sites across modkit pass thresholds (positions covered by *≥* 10 reads in both datasets). Values are given in Supplementary Data 2. See Methods, *Benchmarking of Nanopore inosine calls against Illumina* (A, B) and *Benchmarking against REDIportal* (C).

**Figure S5:**
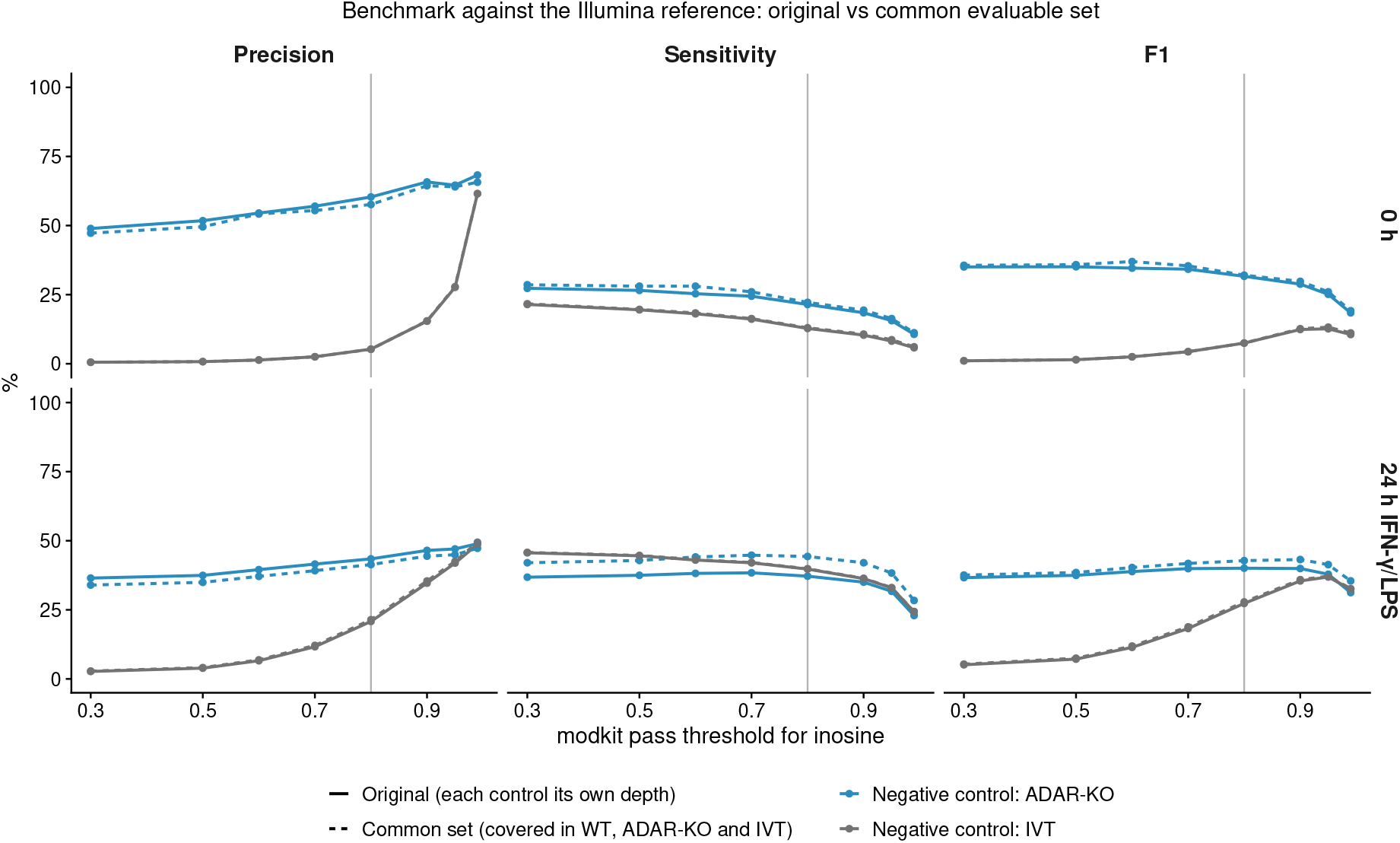
Benchmark of the ADAR1-KO- and IVT-based calls on a common set of sites. Precision, sensitivity and *F*_1_ score of the Nanopore candidate sites relative to the Illumina editing sites across modkit pass thresholds for inosine, with the ADAR1-KO (blue) or IVT (grey) control, in unstimulated cells (0 h, top) and after 24 h of IFN-*γ*/LPS stimulation (bottom). Solid lines: original benchmark, in which each control requires only its own depth (as in Fig. 4C). Dashed lines: Illumina sites and Nanopore calls restricted to the positions covered by at least 10 reads in the Illumina wild-type samples and in the Nanopore WT, ADAR1-KO and IVT samples, so that both controls are evaluated on the same sites (392 Illumina sites at 0 h and 1,099 at 24 h). All other rules as in the main benchmark (frequency *≥* 5%, identical position, one-to-one matching). The vertical line marks the threshold used in the main figures (0.8). Values are given in Supplementary Data 2. See Methods, *Benchmarking of Nanopore inosine calls against Illumina*.

**Figure S6:**
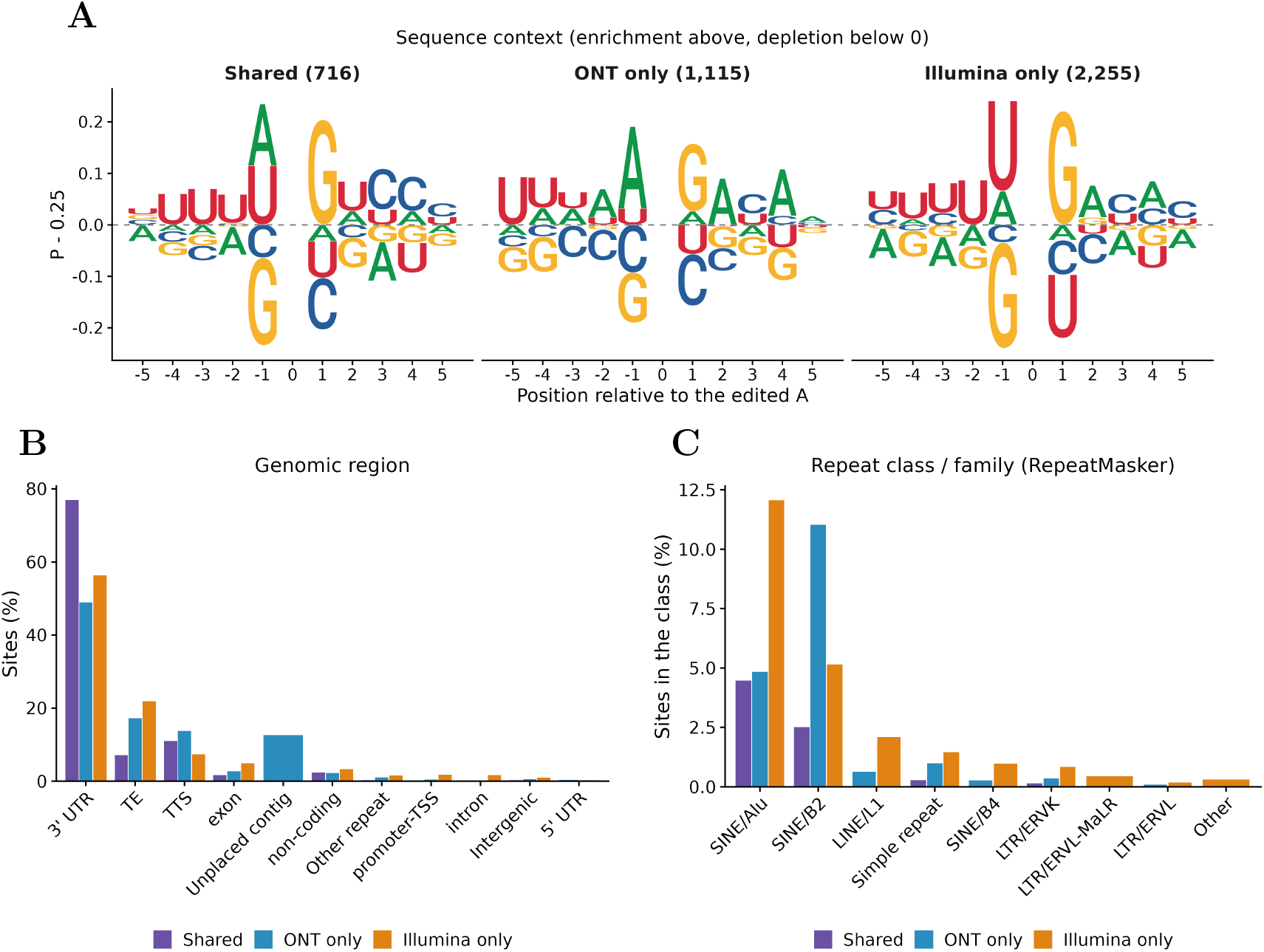
Sequence context and genomic annotation of shared, Nanopore-only and Illumina-only editing sites. Editing sites were classified by identical genomic position as detected by both platforms (Shared, *n* = 716), by Nanopore only (ONT only, *n* = 1,115; ADAR1-KO-based candidate sites, pileup threshold 0.8) or by Illumina only (*n* = 2,255). **(A)** Sequence context of the edited adenosine (*±*5 nt) on the edited strand, shown as the frequency of each base minus 0.25 (the frequency expected at random): bases above the dashed line are enriched, bases below are depleted. The edited adenosine (position 0) is omitted. **(B)** Percentage of sites per genomic region (HOMER annotation; TE, transposable elements (SINE, LINE and LTR); TTS, transcription termination site). Sites on unplaced genomic contigs (`chr*_random`, `chrUn_*`), which are not annotated by HOMER, are shown separately (“Unplaced contig”). **(C)** Percentage of sites in each class that lie in the indicated RepeatMasker repeat class/family (eight most frequent; all others combined as “Other”). Values are given in Supplementary Data 2. See Methods, *Annotation and characterisation of editing sites*.

**Figure S7:**
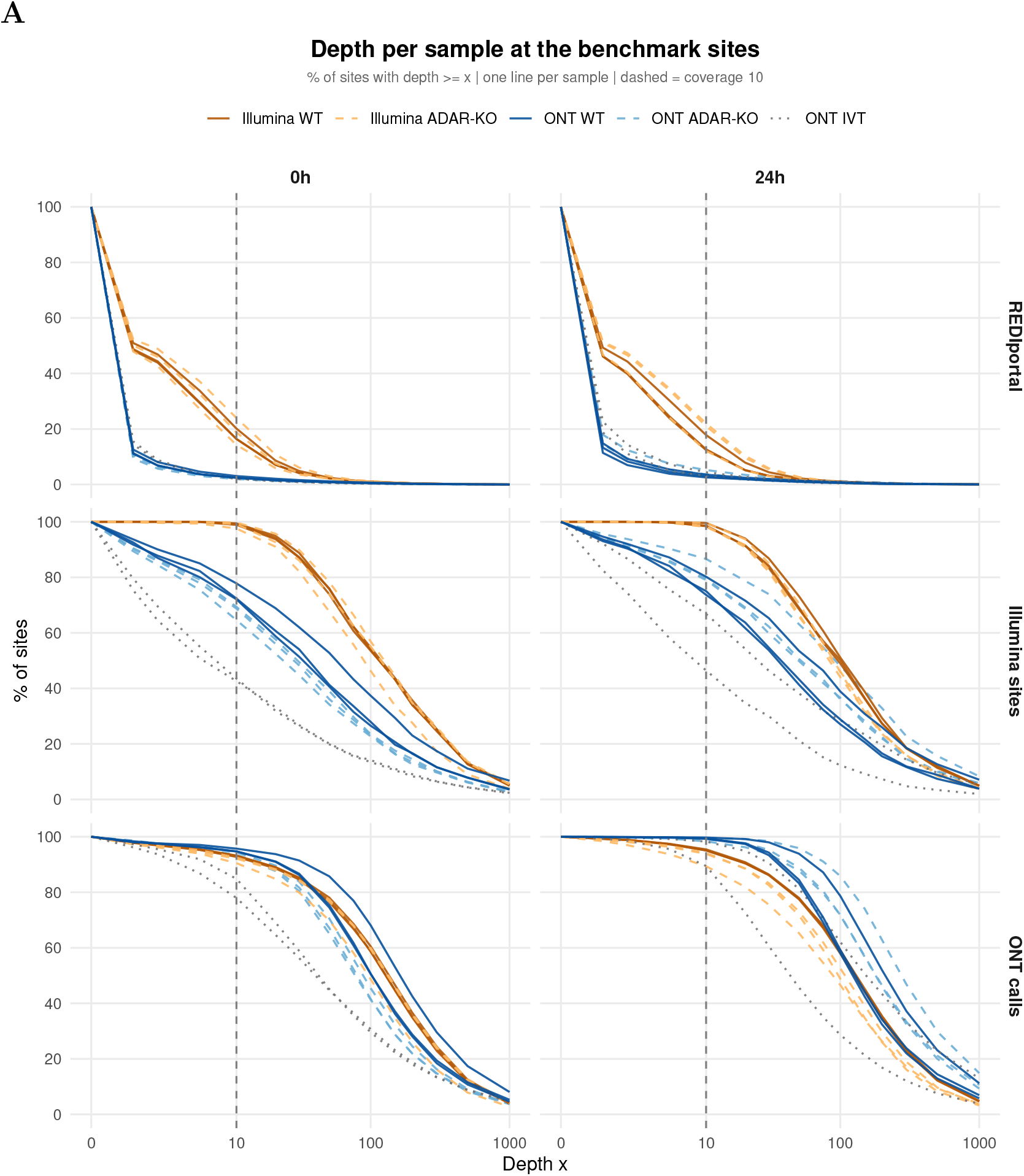
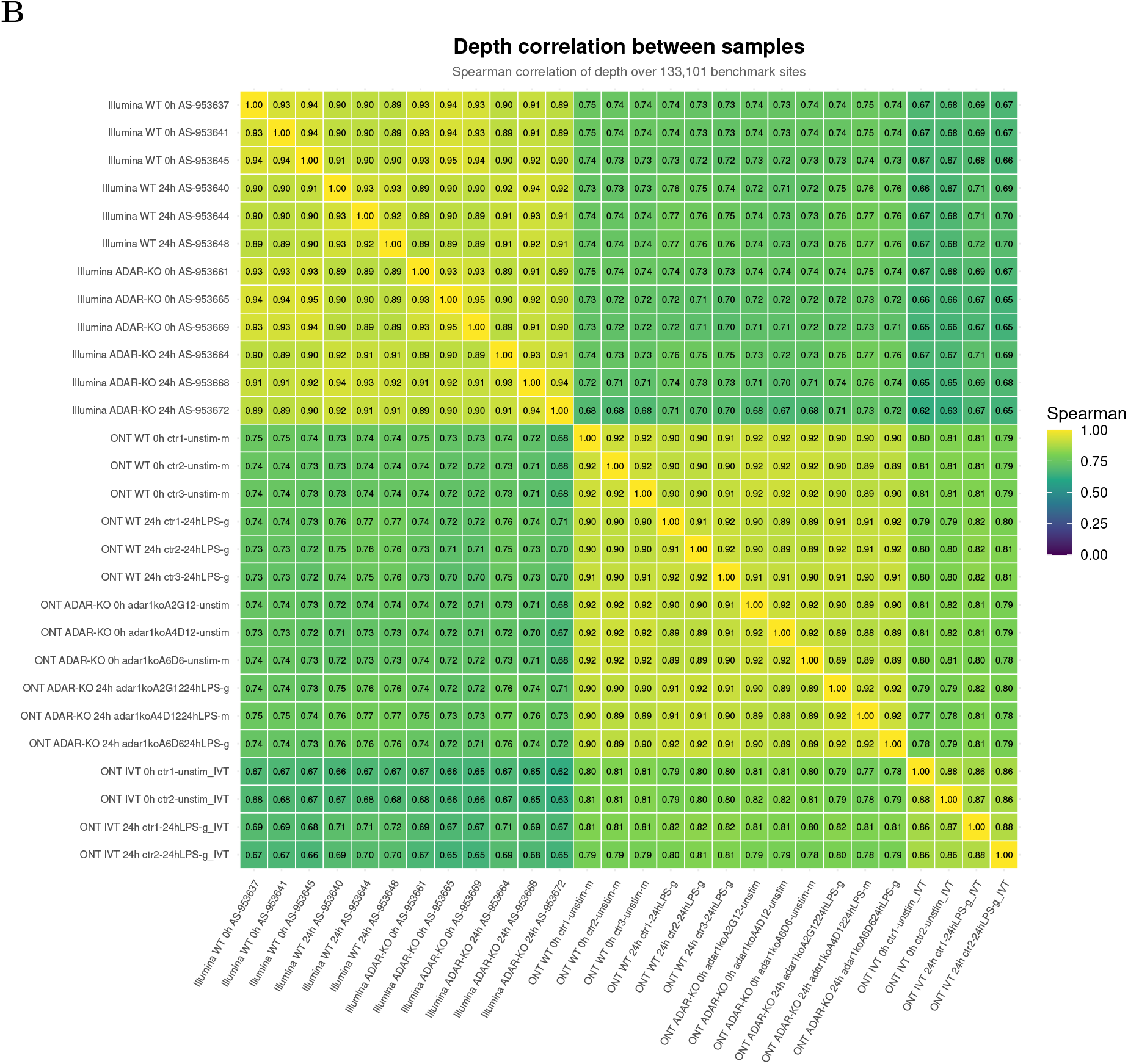
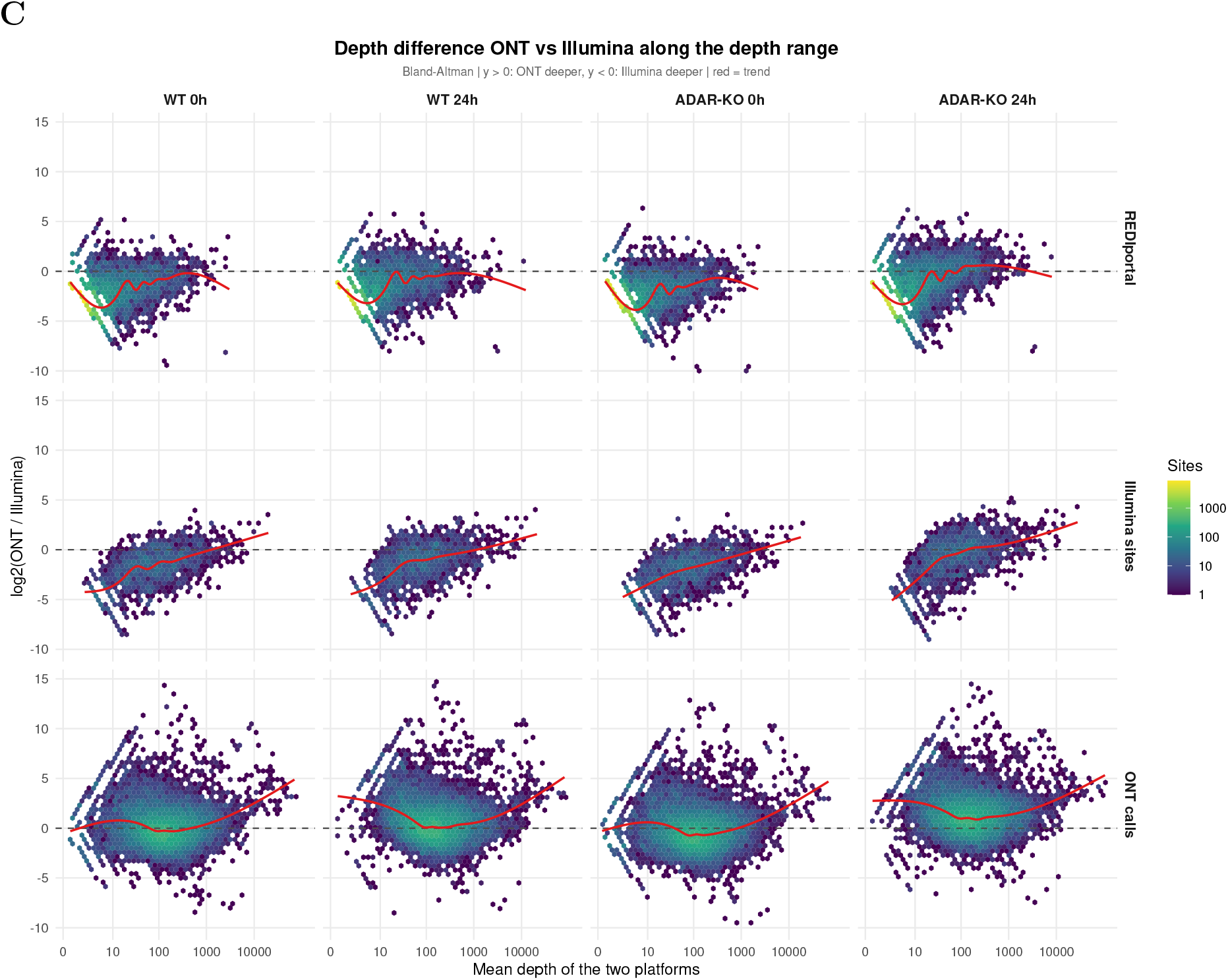
Sequencing depth of the Illumina and Nanopore samples at the benchmarked positions. Read depth was computed with `samtools depth` (MAPQ *≥* 25) at the REDIportal sites, the Illumina editing sites and the Nanopore candidate sites, for each WT, ADAR1-KO and (Nanopore) IVT sample before (0 h) and after 24 h of IFN-*γ*/LPS stimulation. **(A)** Percentage of sites covered by at least the indicated number of reads, one line per sample. **(B)** Spearman correlation of the per-site read depth between all samples. **(C)** log_2_ ratio of Nanopore to Illumina read depth per site (median over replicates) as a function of the mean depth (Bland–Altman); values above 0 indicate higher depth in Nanopore. Red line, smoothed trend (generalised additive model with a cubic regression spline). Values are given in Supplementary Data 2. See Methods, *Read depth at the benchmark sites*.

**Figure S8:**
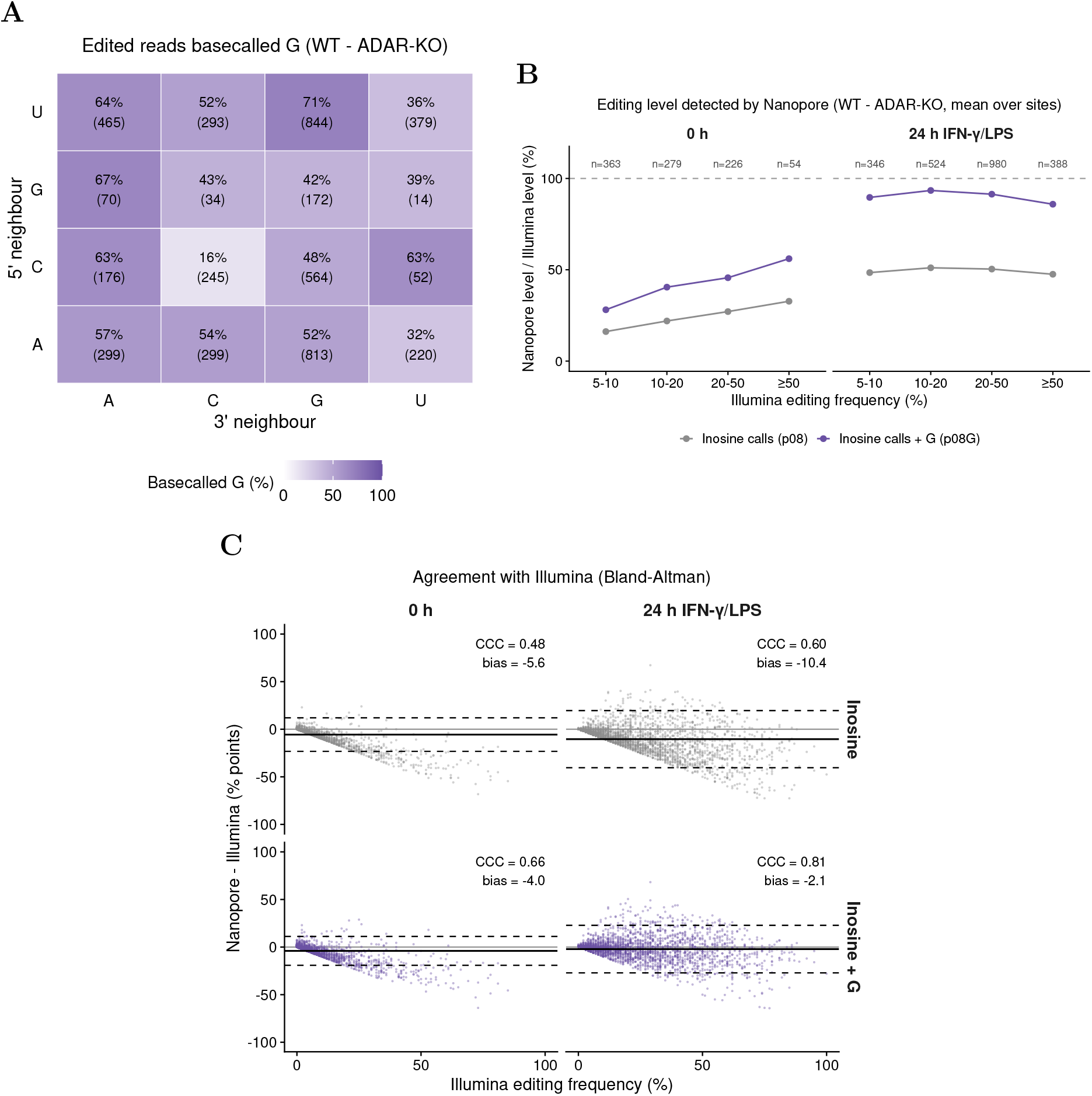
Sequence context, editing level and agreement with Illumina of the inosine reads basecalled as guanosine. **(A)** Percentage of the edited reads basecalled G, by the 5*^′^*and 3*^′^* neighbour of the edited adenosine (RNA strand), at the Illumina editing sites with *≥* 10 Nanopore reads in WT and ADAR1-KO cells (0 h and 24 h pooled). Edited reads are the excess of WT over ADAR1-KO reads (pooled per context), split into reads basecalled G and reads called A with an inosine call (pileup threshold 0.8); numbers in brackets, sites. **(B)** Nanopore editing level (WT minus ADAR1-KO, mean over sites) as a percentage of the Illumina editing frequency (mean over the same sites), from inosine calls only (grey) or from inosine calls plus reads basecalled G (purple), by Illumina editing frequency class; dashed line, equal levels; *n*, sites. **(C)** Agreement of the Nanopore WT level with the Illumina editing frequency at the sites of Fig. 7B (Bland–Altman plot with Illumina as reference): difference between the Nanopore and the Illumina level (percentage points) versus the Illumina frequency, from inosine calls only (top) or with the reads basecalled G (bottom). Solid line, mean difference (bias); dashed lines, limits of agreement (bias *±*1.96 standard deviations); CCC, Lin’s concordance correlation coefficient. Values are given in Supplementary Data 2. See Methods, *Inosine reads basecalled as guanosine*.

## Supplementary Tables

**Table S1:** Samples analysed in this study. RAW 264.7 WT, ADAR1-KO and METTL3-KO cells, unstimulated (0 h) or stimulated for 24 h with IFN-*γ*/LPS, and IVT RNA. Nanopore: direct RNA sequencing (SQK-RNA004), Dorado sup@v6.0; aligned reads are the alignments in the filtered BAM files (mapping quality *≥* 10). Illumina: paired-end RNA-seq; aligned reads are uniquely mapped, deduplicated mates. Median depth and sites with *≥* 10 reads refer to the 2,971 Illumina editing sites (`samtools depth`, mapping quality *≥* 25); not computed for METTL3-KO, which was used only for **Fig. 2**. Figures: main figures in which the sample is used. Raw data: NCBI BioProject PRJNA1232413. See Methods, *Nanopore direct RNA sequencing data processing*, *Illumina RNA-seq and A-to-I editing detection* and *Read depth at the benchmark sites*.

| Platform | Sample | Genotype | Time | Rep. | Aligned reads (M) | Median depth | Sites ≥ 10 reads (%) | Figures |
|---|---|---|---|---|---|---|---|---|
| Nanopore | ctr1-unstim-m | WT | 0 h | 1 | 8.7 | 56 | 77.8 | 2–6 |
| Nanopore | ctr2-unstim-m | WT | 0 h | 2 | 6.7 | 32 | 72.1 | 2–6 |
| Nanopore | ctr3-unstim-m | WT | 0 h | 3 | 5.9 | 34 | 72.4 | 2–6 |
| Nanopore | ctr1-24hLPS-g | WT | 24 h | 1 | 11.3 | 63 | 80.2 | 2–6 |
| Nanopore | ctr2-24hLPS-g | WT | 24 h | 2 | 8.0 | 37 | 75.0 | 2–6 |
| Nanopore | ctr3-24hLPS-g | WT | 24 h | 3 | 6.2 | 38 | 73.7 | 2–6 |
| Nanopore | adar1koA2G12-unstim | ADAR1-KO | 0 h | 1 | 4.9 | 29 | 69.4 | 2–6 |
| Nanopore | adar1koA4D12-unstim | ADAR1-KO | 0 h | 2 | 5.3 | 22 | 64.6 | 2–6 |
| Nanopore | adar1koA6D6-unstim-m | ADAR1-KO | 0 h | 3 | 5.0 | 27 | 68.8 | 2–6 |
| Nanopore | adar1koA2G1224hLPS-g | ADAR1-KO | 24 h | 1 | 9.4 | 52 | 79.5 | 2–6 |
| Nanopore | adar1koA4D1224hLPS-m | ADAR1-KO | 24 h | 2 | 14.0 | 90 | 86.6 | 2–6 |
| Nanopore | adar1koA6D624hLPS-g | ADAR1-KO | 24 h | 3 | 10.6 | 49 | 79.0 | 2–6 |
| Nanopore | mettl3koP1E5-unstim | METTL3-KO | 0 h | 1 | 5.3 | – | – | 2 |
| Nanopore | mettl3koP2F9-unstim | METTL3-KO | 0 h | 2 | 4.9 | – | – | 2 |
| Nanopore | mettl3koP2E2-unstim | METTL3-KO | 0 h | 3 | 5.9 | – | – | 2 |
| Nanopore | mettl3koP1E524hLPS-g | METTL3-KO | 24 h | 1 | 5.4 | – | – | 2 |
| Nanopore | mettl3koP2F924hLPS-g | METTL3-KO | 24 h | 2 | 5.0 | – | – | 2 |
| Nanopore | mettl3koP2E224hLPS-g | METTL3-KO | 24 h | 3 | 4.7 | – | – | 2 |
| Nanopore | ctr1-unstim_IVT | IVT | 0 h | 1 | 11.5 | 5 | 42.6 | 2–6 |
| Nanopore | ctr2-unstim_IVT | IVT | 0 h | 2 | 9.7 | 6 | 43.6 | 2–6 |
| Nanopore | ctr1-24hLPS-g_IVT | IVT | 24 h | 1 | 25.5 | 24 | 66.8 | 2–6 |
| Nanopore | ctr2-24hLPS-g_IVT | IVT | 24 h | 2 | 12.4 | 8 | 46.4 | 2–6 |
| Illumina | AS-953637 | WT | 0 h | 1 | 59.2 | 118 | 99.4 | 1, 4–6 |
| Illumina | AS-953641 | WT | 0 h | 2 | 59.2 | 116 | 98.9 | 1, 4–6 |
| Illumina | AS-953645 | WT | 0 h | 3 | 61.3 | 113 | 99.4 | 1, 4–6 |
| Illumina | AS-953640 | WT | 24 h | 1 | 63.2 | 103 | 99.6 | 1, 4–6 |
| Illumina | AS-953644 | WT | 24 h | 2 | 56.0 | 96 | 98.4 | 1, 4–6 |
| Illumina | AS-953648 | WT | 24 h | 3 | 57.3 | 100 | 98.7 | 1, 4–6 |
| Illumina | AS-953661 | ADAR1-KO | 0 h | 1 | 57.6 | 120 | 99.2 | 1, 4–6 |
| Illumina | AS-953665 | ADAR1-KO | 0 h | 2 | 68.1 | 122 | 99.7 | 1, 4–6 |
| Illumina | AS-953669 | ADAR1-KO | 0 h | 3 | 51.8 | 91 | 97.6 | 1, 4–6 |
| Illumina | AS-953664 | ADAR1-KO | 24 h | 1 | 49.8 | 86 | 98.6 | 1, 4–6 |
| Illumina | AS-953668 | ADAR1-KO | 24 h | 2 | 57.4 | 91 | 99.6 | 1, 4–6 |
| Illumina | AS-953672 | ADAR1-KO | 24 h | 3 | 60.7 | 83 | 98.1 | 1, 4–6 |

**Table S2:** Sites overlapping known SNPs. Number of editing sites at the position of a single-nucleotide polymorphism of the Mouse Genomes Project (v5, 36 strains): at any SNP, at a SNP with a dbSNP 142 rs ID, or at a SNP carried by BALB/cJ, the genetic background of RAW 264.7. Percentages are relative to the total number of sites in the set. Nanopore sets: candidate sites at pileup threshold 0.8; site classes (Shared, ONT only, Illumina only) by identical position, ADAR1-KO-based. Related to **Fig. S1A**; see Methods, *Annotation and characterisation of editing sites*.

| Site set | Sites | Sites at a known SNP |  |  |
|---|---|---|---|---|
|  |  | any strain | with dbSNP 142 rs ID | carried by BALB/cJ |
| Illumina | 2,971 | 96 (3.2%) | 72 | 0 |
| ONT (ADAR1-KO) | 1,831 | 54 (2.9%) | 43 | 0 |
| ONT (IVT) | 2,251 | 44 (2.0%) | 34 | 0 |
| Shared | 716 | 28 (3.9%) | 24 | 0 |
| ONT only | 1,115 | 26 (2.3%) | 19 | 0 |
| Illumina only | 2,255 | 68 (3.0%) | 48 | 0 |

**Table S3:** Benchmark at 24 h with an Illumina reference extended by stimulation-specific editing sites. Nanopore candidate sites (ADAR1-KO-based, pileup threshold 0.8) scored against three Illumina references at 24 h: the reference of the main benchmark (sites edited at both time points), the reference extended by the positions with A-to-G substitutions at 24 h only, and these positions alone. Same rules as the main benchmark (coverage *≥* 10 in both platforms and in the Nanopore negative control, frequency *≥* 5%, identical position, one-to-one matching). TP, true positives; FP, false positives; FN, false negatives. *^a^*Only Nanopore calls not contained in the original reference were scored. The 24 h-only positions rest on two or three replicates of a single time point and were not used as reference in the main analysis. See Methods, *Stimulation-specific editing sites*.

| Illumina reference (24 h) | Reference sites | Nanopore calls | TP | FP | FN | Sensitivity (%) | Precision (%) | *F*<sub>1</sub> (%) |
|---|---|---|---|---|---|---|---|---|
| Original (both time points) | 1,896 | 1,623 | 705 | 918 | 1,191 | 37.2 | 43.4 | 40.1 |
| Extended (+ 24 h-only positions) | 3,998 | 1,623 | 889 | 734 | 3,109 | 22.2 | 54.8 | 31.6 |
| 24 h-only positions alone<sup>*a*</sup> | 2,102 | 918 | 184 | 734 | 1,918 | 8.8 | 20.0 | 12.2 |

**Table S4:** Benchmark of the Nanopore candidate sites against the Illumina editing sites, including the Jaccard index. Nanopore candidate sites (pileup threshold 0.8) obtained with the ADAR1-KO or IVT control, from inosine calls only or with the reads basecalled G counted as modified (inosine + G), scored against the Illumina editing sites of the same time point: coverage *≥* 10 in both platforms and in the Nanopore negative control, frequency *≥* 5%, identical position, one-to-one matching. TP, true positives (reference sites called by Nanopore); FP, false positives (Nanopore calls not in the Illumina reference); FN, false negatives (reference sites not called by Nanopore). Because the Illumina sites are a reference and not a ground truth, FP and FN are not necessarily errors. Sensitivity, TP/(TP + FN); precision, TP/(TP + FP); *F*_1_, harmonic mean of both; Jaccard index, TP/(TP + FP + FN), the fraction of all sites detected by either platform that are detected by both. The rows with inosine calls only correspond to **Fig. 4C** at threshold 0.8, the comparison of both call sets to **Fig. 7C**. Values are also given in Supplementary Data 2. See Methods, *Benchmarking of Nanopore inosine calls against Illumina* and *Inosine reads basecalled as guanosine*.

| Control | Calls | Illumina sites | Nanopore calls | TP | FP | FN | Sensitivity (%) | Precision (%) | *F*<sub>1</sub> (%) | Jaccard (%) |
|---|---|---|---|---|---|---|---|---|---|---|
| *0 h* |  |  |  |  |  |  |  |  |  |  |
| ADAR1-KO | inosine | 667 | 237 | 143 | 94 | 524 | 21.4 | 60.3 | 31.6 | 18.8 |
| ADAR1-KO | inosine + G | 667 | 445 | 265 | 180 | 402 | 39.7 | 59.6 | 47.7 | 31.3 |
| IVT | inosine | 415 | 1,004 | 53 | 951 | 362 | 12.8 | 5.3 | 7.5 | 3.9 |
| IVT | inosine + G | 415 | 1,060 | 2 | 1,058 | 413 | 0.5 | 0.2 | 0.3 | 0.1 |
| *24 h IFN-γ/LPS* |  |  |  |  |  |  |  |  |  |  |
| ADAR1-KO | inosine | 1,896 | 1,623 | 705 | 918 | 1,191 | 37.2 | 43.4 | 40.1 | 25.1 |
| ADAR1-KO | inosine + G | 1,896 | 3,033 | 1,168 | 1,865 | 728 | 61.6 | 38.5 | 47.4 | 31.1 |
| IVT | inosine | 1,102 | 2,105 | 438 | 1,667 | 664 | 39.7 | 20.8 | 27.3 | 15.8 |
| IVT | inosine + G | 1,102 | 1,182 | 13 | 1,169 | 1,089 | 1.2 | 1.1 | 1.1 | 0.6 |

